# Sarcomeres regulate cardiomyocyte maturation through MRTF-SRF signaling

**DOI:** 10.1101/824185

**Authors:** Yuxuan Guo, Blake D. Jardin, Isha Sethi, Qing Ma, Behzad Moghadaszadeh, Emily C. Troiano, Michael A. Trembley, Eric M. Small, Guo-Cheng Yuan, Alan H. Beggs, William T. Pu

**Affiliations:** Department of Cardiology, Boston Children’s Hospital, 300 Longwood Ave, Boston, MA 02115, USA; Department of Biostatistics and Computational Biology, Dana-Farber Cancer Institute, 450 Brookline Ave, Boston, MA 02215, USA; Division of Genetics and Genomics, The Manton Center for Orphan Disease Research, Boston Children’s Hospital and Harvard Medical School, Boston, MA 02115, USA; University of Rochester School of Medicine and Dentistry, Rochester, NY 14642, USA; Harvard Stem Cell Institute, 7 Divinity Avenue, Cambridge, MA 02138, USA

## Abstract

Cardiomyocyte maturation is essential for robust heart contraction throughout life. The signaling networks governing cardiomyocyte maturation remain poorly defined. Our prior studies established the transcription factor SRF as a key regulator of the assembly of sarcomeres, the contractile unit of cardiomyocytes. Whether sarcomeres regulate other aspects of maturation remains unclear. Here we generated mice with cardiomyocyte specific, mosaic mutation of α-actinin-2 (*Actn2*), a key organizer of sarcomeres, to study its cell-autonomous role in cardiomyocyte maturation. In addition to the expected structural defects, *Actn2* mutation triggered dramatic transcriptional dysregulation, which strongly correlated with transcriptional changes observed in SRF-depleted cardiomyocytes. *Actn2* mutation increased monomeric actin, which perturbed the nuclear localization of the SRF cofactor MRTFA. Overexpression of a dominant-negative MRTFA mutant was sufficient to recapitulate the transcriptional and morphological defects in *Actn2* and *Srf* mutant cardiomyocytes. Together, we demonstrate that ACTN2-based sarcomere assembly and MRTF-SRF signaling establish a positive feedback loop that promotes cardiomyocyte maturation.

## Introduction

One of the final steps in the development of the mature heart is cardiomyocyte (CM) maturation. This process is essential for the generation of robust CMs that can sustain forceful contractions throughout postnatal life. The transcriptional, morphological, and functional hallmarks of mature CMs have been well described (*1*), but little is known about how CMs acquire these features during development. This knowledge gap obscures the contribution of CM maturation to both the pathogenesis of inherited cardiomyopathies and the loss of heart regenerative capacity. It also impairs efforts to engineer mature stem cell-derived cardiac tissues for cell therapy, in vitro disease modeling, and pharmacological testing.

The transcriptional and ultrastructural changes that occur during CM maturation are likely coordinated. For example, the maturation of sarcomeres, the contractile apparatus of CMs, requires proper transcription and isoform switching of genes encoding sarcomeric components. The products of sarcomere gene expression require efficient assembly into expanding myofibrils. Similarly, the maturation of CM electrophysiology requires proper ion channel expression and localization to specialized ultrastructural hallmarks of adult CMs. These structures include the transverse tubules (T-tubules), a network of plasma membrane invaginations, and dyads, formed by the close apposition of T-tubules to the junctional sarcoplasmic reticulum. T-tubules and dyads facilitate and coordinate cardiomyocyte excitation-contraction coupling. CM metabolic maturation is characterized by extensive upregulation of genes that regulate fatty acid catabolism and oxidative phosphorylation. Meanwhile, mitochondria grow dramatically in both number and size, providing an extensive inner membrane and cristae network that is essential for efficient oxidative energy production.

Mechanisms responsible for the mutual communication between ultrastructural components and transcriptional regulators in CM maturation are beginning to emerge. We recently reported that the transcription factor serum response factor (SRF) is essential for CM maturation (*2*). SRF directly activates the transcription of genes regulating sarcomere assembly, electrophysiology, and mitochondrial metabolism. This subsequently drives the proper morphogenesis of mature ultrastructural features of myofibrils, T-tubules, and mitochondria. Sarcomeres are essential mediators that translate transcriptional information into ultrastructural changes, partly by providing structural support for the biogenesis of T-tubules and mitochondria. Accumulated evidence has implicated that sarcomeres can function as a regulator of signal transduction and transcription (*3*, *4*). However, the role of sarcomere signaling in CM maturation and its molecular mechanism are yet to be determined.

In non-myocytes, SRF is influenced by the polymeric state of actin through SRF transcriptional cofactors myocardin-related transcription factor A and B (MRTFA/B; also known as MKL1/2). Monomeric actin (G-actin) interaction with MRTFs leads to retention of MRTFs in the cytoplasm and inhibition of its activity as a transcriptional co-activator (*5*). G-actin polymerization to form polymeric, filamentous actin (F-actin) decreases G-actin concentration, derepresses MRTFs, and activates SRF. In CMs, F-actin is a major constituent of sarcomere thin filaments (*6*). MRTFA/B depletion in CMs triggers severe cardiomyopathy shortly after birth, a critical time window when major CM maturation events occur (*7*). Thus we hypothesized that sarcomere assembly in CMs reduces G-actin concentration, which then activates MRTF-SRF signaling required for CM maturation.

Investigating the role of sarcomeres is technically challenging. In vitro cell culture is not optimal for these studies because mature CMs cannot be maintained in cell culture and their sarcomeres rapidly become disorganized. Standard organ-wide genetic manipulations of sarcomere genes in mice often cause cardiac decompensation or lethality, which complicate the analysis of sarcomeres in CM maturation. To circumvent these issues, we recently established cardiac genetic mosaic analysis to dissect CM maturation. In this experimental paradigm, we use low dose adeno-associated virus to genetically manipulate a low fraction (10-20%) of CMs via cDNA overexpression or Cas9 or Cre-loxP conditional mutagenesis (*8*, *9*). This approach circumvents lethality and the secondary effects of heart dysfunction, enabling us to study the cell-autonomous function of sarcomere genes in CM maturation within an overall normal physiological context.

α-actinin-2 (ACTN2) is a key component of sarcomeric Z-lines and is essential for sarcomere assembly (*10*). In this study, we used both Cas9-mediated somatic mutagenesis and a conditional hypomorphic *Actn2* allele to generate hearts with mosaic *Actn2* mutation. These experiments revealed that ACTN2-dependent sarcomere assembly cell-autonomously regulates the transcriptional program required for CM maturation through MRTF-SRF signaling.

## Results

### Cas9-based somatic mutagenesis of *Actn2* in CM maturation

We previously established a CRISPR/Cas9-AAV9-based somatic mutagenesis (CASAAV) system (Fig. S1A) to determine the role of a given gene in CM maturation (*8*, *11*). We produced two AAV vectors targeting *Actn2*. Each AAV expresses two independent gRNAs and Cre recombinase under the control of the CM-specific cardiac troponin T (cTnT) promoter (*12*, *13*). These AAVs were injected into postnatal day one (P1) mice carrying a Cre-activatable Cas9-2A-GFP allele (*14*) to trigger Cas9-based mutagenesis of *Actn2*. We verified successful depletion of ACTN2 protein in ~60% of transduced (GFP+) CMs at P30 by immunostaining (Fig. S1B-C). Mutant CMs exhibited defective T-tubulation and maturational CM hypertrophy (Fig. S1D-F), indicating CM maturation defects. These data encouraged us to mechanistically analyze the link between ACTN2 and CM maturation.

### Generation of a conditional hypomorphic allele of *Actn2*

To facilitate more detailed studies, we generated a floxed *Actn2* allele (*Actn2*^*F*^), in which the genomic region containing exons 2, 3, and 4 was flanked by LoxP sequences (Fig. 1A). Heterozygous (*Actn2*^*F/+*^) and homozygous (*Actn2*^*F/F*^) mice were fertile and exhibited normal heart function prior to Cre-mediated recombination (Fig. S2A-C).

**Fig. 1.**
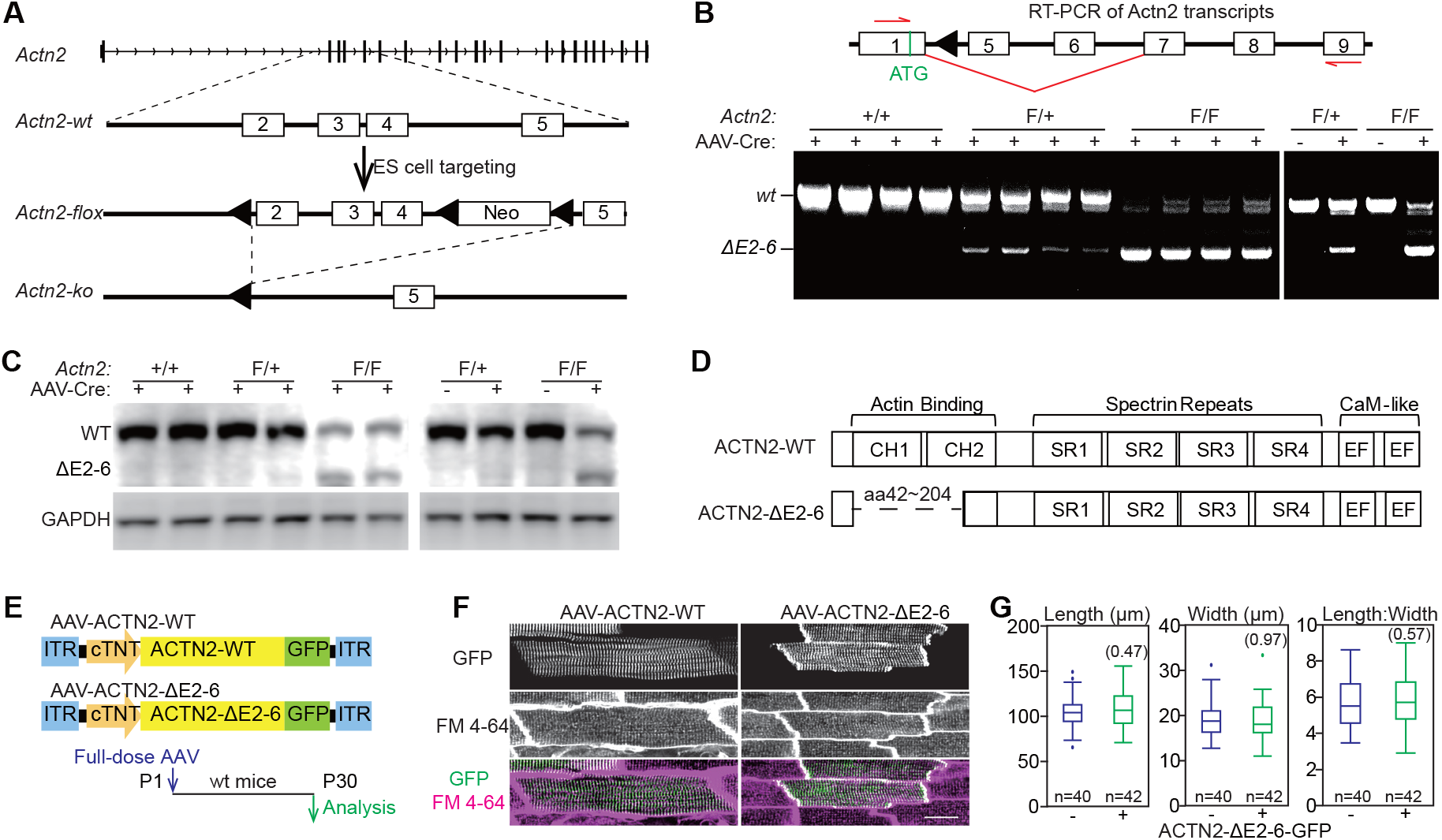
Generation of a conditional hypomorphic allele of *Actn2*. **(A)** Design of the *Actn2*^*F*^ allele. **(B)** RT-PCR identified ectopic splicing between exon 1 and 7 following Cre-LoxP recombination. **(C)** ACTN2 western blot of control and mutant heart extracts using a C-terminal ACTN2 antibody. In **(B)** and **(C)**, a high dose of AAV-Cre was injected into P1 pups and ventricles were analyzed at P7. **(D)** Domain structure of wild-type and mutant ACTN2 proteins. **(E)** Experimental design of AAV-mediated overexpression of wild-type and mutant ACTN2 tagged with GFP in CMs. **(F)** *In situ* imaging of GFP and T-tubules in the heart. Scale bar, 20 μm. **(G)** Morphometric measurement of isolated CMs from hearts with mosaic overexpression of mutant ACTN2. In box plots, the horizontal line within the box represents the median; the ends of the box represent the 25th and 75th quantiles; whiskers extend 1.5 times the interquartile range from the 25th and 75th percentiles; dots represent possible outliers. Two-tailed student’s *t*-test: Non-significant P-values are labeled in parentheses.

To determine if Cre-mediated recombination of *Actn2*^*F*^ in maturing CMs caused *Actn2* inactivation, we treated P1 Actn2^F/F^ or control pups with AAV-Cre, an AAV vector expressing Cre driven by the cardiomyocyte-selective cTnT promoter (*12*). At P7, we analyzed *Actn2* transcripts by reverse transcriptase PCR (RT-PCR) using primers that anneal to exons 1 and 9. The size of the major RT-PCR product in Cre-recombined samples was not consistent with simple deletion of exons 2-4 (Fig. 1B). DNA sequencing showed that the major RT-PCR amplicon in Cre-recombined samples resulted from ectopic splicing between exons 1 and 7 (Fig. S3A), which we denote as *Actn2*^*ΔE2-6*^.

The exon 1-7 junction preserves the open reading frame and is expected to produce a mutant ACTN2 protein. To detect this protein, we treated P1 hearts with 1.1×10^10^ vg/g AAV-Cre, a dose calibrated to transduce more than 75% of CMs. At P7, we confirmed the expression of ACTN2^ΔE2-6^ by western blot using an ACTN2 antibody that recognizes the C-terminus of ACTN2 (Fig. 1C). *Actn2*^*ΔE2-6*^ transcript and protein were detected only when both the floxed allele and Cre were present, indicating that they resulted from Cre-mediated recombination and not from genetic modification of *Actn2* or an effect of AAV-Cre alone.

ACTN2^ΔE2-6^ lacks amino acids 42-204 (Fig. 1D), encompassing the entire calponin homology 1 (CH1) domain and the N-terminal half of the CH2 domain, the actin-binding domains of ACTN2. We treated mice with AAV expressing ACTN2^ΔE2-6^-GFP to determine its impact on CM maturation, using an AAV expressing wild-type (WT) ACTN2-GFP (ACTN2^WT^) as control (Fig. 1E). Optical confocal sectioning of hearts perfused with the membrane stain FM4-64 (*15*) showed that ACTN2^ΔE2-6^-GFP exhibited the same striated pattern as ACTN2^WT^-GFP, consistent with Z-line localization (Fig. 1F). In addition, ACTN2^ΔE2-6^-GFP displayed greater localization on intercalated discs (Fig. 1F), suggesting an actin-independent interaction between ACTN2 and the intercalated disc. No defects in sarcomere organization (Fig. 1F), T-tubule morphology (Fig. 1F), or cell size and shape (Fig. 1G) were detected in ACTN2^ΔE2-6^-GFP expressing CMs. These data indicate that ACTN2^ΔE2-6^-GFP was not deleterious to these readouts of CM maturation. Consistent with these data, echocardiography showed no functional defects when ~75% of CMs were transduced with AAV-ACTN2^ΔE2-6^-GFP (Fig. S2D-F). Thus, ACTN2^ΔE2-6^ is a hypomorphic allele that is unlikely to exert a dominant-negative impact on CM maturation.

### Mosaic *Actn2* mutation in the heart

*Actn2*^*F*^ alleles were combined with Cre-inducible fluorescent protein (FP) reporter alleles *Rosa*^*Cas9GFP*^ *(14)* or *Rosa*^*tdTomato*^ *(16)*, so that FP reporters could be used as surrogate markers of AAV-Cre transduced CMs. Full (1.1×10^10^ vg/g)- and high (3.6×10^9^ vg/g)-dose AAV-Cre treatment of P1 *Actn2*^*F/F*^ pups resulted in death within the first two weeks of life (Fig. 2A). High-dose and Mid-dose (1.1×10^9^ vg/g) AAV-Cre triggered acute dilated cardiomyopathy, which was characterized by diminished left ventricular systolic function and dilatation by echocardiography (Fig. 2B). Picro sirius red and wheat germ agglutinin (WGA) staining revealed cardiac fibrosis in the high- and mid-dose AAV-Cre treated hearts (Fig. 2C). Real-time quantitative PCR (RTq-PCR) of heart RNA detected the upregulation of cardiac stress markers *Nppa* and *Nppb* in the groups with cardiac dysfunction (Fig. 2D). These data confirmed a critical role of ACTN2 in heart function.

**Fig. 2.**
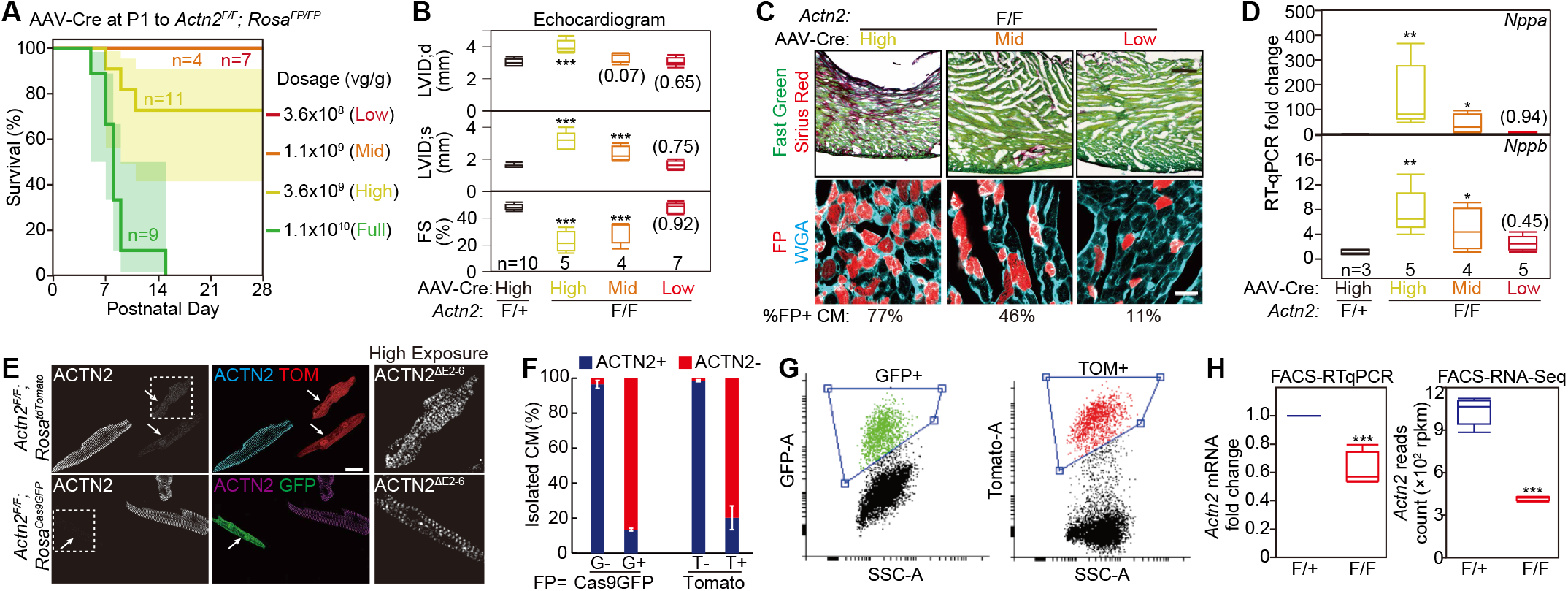
Characterization of mice with mosaic *Actn2* mutation. **(A)** Survival curve of *Actn2*^*F/F*^;*Rosa*^*FP/FP*^ mice treated with low, mid, high, and full doses of AAV-Cre. **(B)** Effect of AAV-Cre dosage on heart function and chamber size at P30. Left ventricle (LV) function and size were assessed echocardiographically by measuring the LV fractional shortening (FS) and internal diameter at end systole (LVID;s) and diastole (LVID;d). **(C)** Effect of AAV-Cre dosage on myocardial fibrosis. P30 heart sections were stained with sirius red/fast green (top; scale bar = 200 μm) or wheat germ agglutinin (WGA, bottom; scale bar = 20 μm). The fraction of FP+ cells is indicated below images. **(D)** RT-qPCR analysis of cardiac stress marker mRNA from P30 heart ventricles. **(E)** Representative images of ACTN2 immunofluorescence in P30 FP+ and FP- CMs that were isolated from the same heart. Arrows point to ACTN2 mutant CMs. Boxed area is enlarged and overexposed to show the presence of mutant ACTN2 protein. **(F)** Quantification of CMs with decreased ACTN2 signal (ACTN2-) among FP+ and FP-CMs. G, Cas9GFP; T, tdTomato. Error bar, SD. n=3 hearts per group **(G)** Representative FACS plots and gating to sort FP+ P14 CMs. **(H)** Measurement of *Actn2* mRNA level in FACS-sorted FP+ CMs by RT-qPCR (left) and RNA-seq (right) at P14. n=5 hearts per group. Box plots, statistical analysis and scale bars are the same as in **Fig. 1**. Two-tailed student’s t-test versus control: *P<0.05, **P<0.01, ***P<0.001. Numbers in parentheses indicate non-significant P-values.

Cardiac dysfunction triggers cascades of secondary effects that confound study of CM maturation (*8*, *9*). To circumvent this problem, we identified a low dose (3.6×10^8^ vg/g) of AAV-Cre that recombined ~11% of CMs without causing overall heart dysfunction, fibrosis, or activation of cardiac stress markers (Fig. 2A-D). We treated *Actn2*^*F/F*^*; Rosa*^*FP/FP*^ mice with this dose of AAV-Cre at P1 and isolated CMs at P30. Immunofluorescent staining of ACTN2 in dissociated CMs (Fig. 2E) showed a dramatic decrease in ACTN2 signal in 80-85% of FP+ CMs (Fig. 2F). However, a low level of ACTN2 immunoreactivity persisted (Fig. 2E, right “High Exposure”), consistent with low residual expression of ACTN2^ΔE2-6^. On the other hand, over 95% of FP-CMs retained strong ACTN2 staining (Fig. 2F). These data showed that ACTN2^ΔE2-6^ expression level was much lower than ACTN2^WT^, further arguing against a dominant-negative effect of ACTN2^ΔE2-6^. We also purified FP+ CMs from AAV-Cre-treated *Actn2*^*F/F*^ and *Actn2*^*F/+*^ mice by flow cytometry (Fig. 2G) and performed RT-qPCR and RNA-Seq to validate the decrease of *Actn2* mRNA in mutant CMs (Fig. 2H). RNA-Seq analysis confirmed aberrant splicing between exons 1 and 7 (Fig. S3B) and dramatically decreased expression of exon 2-6 in fluorescence-activated cell sorting (FACS)-purified mutant CMs (Fig. S3C). Read counts of all other exons were also reduced. Thus, Cre-mediated *Actn2* recombination perturbed overall *Actn2* transcript levels in addition to the skipping of exons 2-6. Residual RNA-Seq signal on exons 2-6 in mutant CMs was likely a consequence of imperfect FACS sorting. Together, these data indicate that FP expression is a reliable surrogate marker of CMs with impaired *Actn2* expression. Furthermore, the data show that we have generated a genetic mosaic model of *Actn2* mutation that allows investigation of the cell-autonomous roles of ACTN2. FP- *Actn2*^*F/F*^ CMs, derived from the same heart as FP+ mutant CMs, could be used as control cells. Alternatively, FP+ CMs from *Actn2*^*F/+*^ littermates treated with AAV-Cre as controls.

### Impact of *Actn2* on morphological maturation

F-actin is a major constituent of sarcomeric thin filaments. To determine the role of ACTN2 in sarcomere organization, we stained dissociated CMs for ACTN2 and phalloidin, which selectively binds F-actin. Compared to ACTN2+ CMs, ACTN2 mutant CMs had drastically reduced F-actin, as measured by phalloidin fluorescence intensity (Fig. 3A-B). Scattered F-actin myofibrils were retained in the mutant CMs and were associated with the plasma membrane, while short fragments of F-actin (Fig. 3A, arrows) could be observed in the interior of mutant CMs. To further study the effect of *Actn2* mutation on sarcomere ultrastructure, we performed electron microscopy (EM) on CMs that were purified by FACS (FACS-EM) (*2*, *15*). Z-lines, the electron-dense boundaries of sarcomeres organized by ACTN2, were noticeably diminished in the remnant myofibrils in mutant CMs (Fig. 3C, red arrows). Longitudinal sarcomere filaments were disorganized and M-lines, an ultrastructural hallmark of mature sarcomeres (*17*), were not present in mutant CMs (Fig. 3C, yellow arrows). These data confirmed the role of ACTN2 as a key organizer of sarcomeres.

**Fig. 3.**
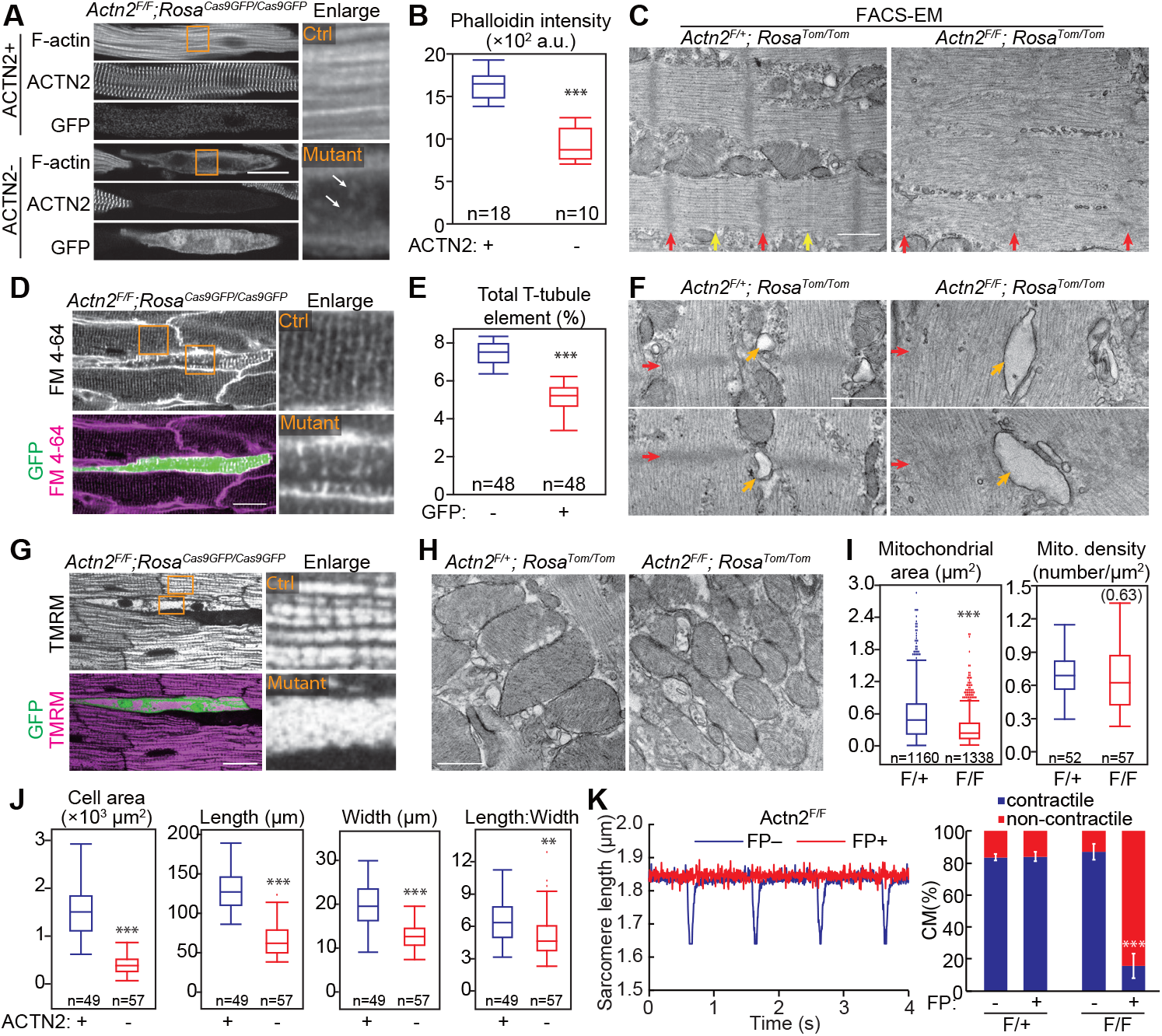
The impact of *Actn2* mutation on CM morphological maturation. Mosaic mutation of *Actn2* was achieved by low dose AAV-Cre treatment of *Actn2*^*F/F*^*; Rosa^FP/FP^* mice. Control cells were either FP- or ACTN2+ cells from the same heart **(A,D,G,J,K)** or FP+ CMs from *Actn2*^*F/+*^*; Rosa^FP/FP^* mice **(C,F,H)**. Mice were treated on P1 and analyzed at P30. **(A)** Phalloidin staining on control and *Actn2* mutant CMs. Arrows point to small F-actin fragments. **(B)** Quantification of fluorescence intensity of phalloidin staining. **(C)** EM analysis of sarcomere ultrastructure in FACS-sorted Tomato+ control and mutant CMs. Red arrows point to Z-lines, and yellow arrows to M-lines. **(D)** Confocal optical sections of hearts perfused with plasma membrane dye FM 4-64. Stained membranes within CMs are T-tubules. **(E)** T-tubule quantification by AutoTT. **(F)** EM analysis of T-tubules in FACS-sorted Tomato+ control and mutant CMs. Red arrows indicate Z-lines, and orange arrows indicate T-tubules. **(G)** Confocal optical sections of hearts perfused with mitochondrial dye TMRM. **(H)** EM analysis of mitochondria. **(I)** Quantification of mitochondrial area and density by EM. **(J)** CM geometry, measured on dissociated CMs. **(K)** Representative traces (left) and quantification (right) of contractile CMs upon electrical stimulation. Error bar is SD. n=3 hearts per group. In **(A,D,G)**, boxed areas are enlarged to show detailed structures, and scale bars are 20 μm. In **(C,F,H)**, scale bars are 500 nm. In **(B,E,I,J)**, box plots were generated similar to **Fig. 1**. All statistical analyses are the same as **Fig. 2**.

We next investigated the impact of *Actn2* mutation on other structural hallmarks of CM maturation, using confocal optical sectioning of hearts perfused with the plasma membrane dye FM 4-64 (*15*). This allowed us to assess the cell-autonomous effect of *Actn2* mutation on T-tubulation. Mutant, FP+ CMs exhibited reduced T-tubule content and organization, which was confirmed by quantification using AutoTT software (*18*) (Fig. 3D-E). Mutant CMs exhibited disrupted T-tubule organization, and residual T-tubules had greater FM4-64 signal intensity (Fig. 3D). FACS-EM analysis showed that mutant CMs contained abnormal, dramatically dilated T-tubules, which often remained adjacent to remnant sarcomere Z-lines (Fig. 3F).

To study the effect of *Actn2* mutation on mitochondrial organization, we performed confocal optical sectioning of hearts perfused with the mitochondrial dye tetramethylrhodamine methyl ester (TMRM), which indicates mitochondrial membrane potential and organization. Overall TMRM intensity was retained, indicating preservation of mitochondrial membrane potential. However, the mitochondria showed dramatic disorganization in *Actn2* mutant CMs (Fig. 3G and Fig. S4). FACS-EM revealed decreased mitochondrial size, consistent with CM immaturity (Fig. 3H-I). However, mitochondrial density did not change, suggesting that mitochondrial number increased to compensate for reduced size.

To assess the effect of *Actn2* mutation on maturational CM growth, we measured the geometry of immunostained CMs isolated from AAV-Cre-treated *Actn2*^*F/F*^*;Rosa*^*FP/FP*^ hearts. Mutant CMs (FP+) had decreased cell area due to reduction of both cell length and width (Fig. 2E and Fig. 3J). The length:width ratio also decreased, indicating that ACTN2 plays a more profound role in CM elongation than widening during maturation. We also determined if *Actn2* mutation affected CM contractility by measuring the length of sarcomeres in paced, dissociated CMs. Cell contraction was completely abolished in ~80% of FP+ *Actn2*^*F/F*^ CMs (Fig. 3K), which was the same as the frequency of FP+ CMs with mutant ACTN2 (Fig. 2F).

Together, these data indicate that ACTN2 is critical for normal structural maturation of CMs.

### Impact of *Actn2* on transcription through MRTF-SRF signaling

We hypothesized that ACTN2 regulates signal transduction and gene transcription, beyond its canonical role as a structural protein. To test this hypothesis, we injected AAV-Cre at P1, purified control and mutant CMs by FACS at P14, and performed RNA-Seq analysis using a recently established protocol (*2*) (FACS-RNA-seq, Fig. 4A). Five biological replicates for each group were clearly separated by principal component analysis (Fig. 4B; Suppl. Table 1). Differential expression analysis identified 1352 and 1287 genes that were up- or down-regulated in mutant CMs respectively, using Padj<0.05 as the cutoff (Fig. 4C; Suppl. Data 1). Since SRF is a key transcription factor regulating CM maturation (*2*), we compared *Actn2* mutant and *Srf* knockout (KO) RNA-seq data (*2*) produced using the same protocol. 60.0% of down-regulated genes in *Srf* KO CMs were also down-regulated in *Actn2* mutant CMs, and 57.1% of up-regulated genes in *Srf* KO CMs were also up-regulated in *Actn2* mutant CMs (Fig. 4D). The fold-changes of these shared genes were highly correlated (r=0.93) between *Actn2* mutant and *Srf* KO datasets (Fig. 4D). Gene set enrichment analysis (GSEA) (*19*) showed that the top 10 down-regulated gene ontology (GO) terms, which were related to metabolism, mitochondria, and translation, were identical in the two experiments (Fig. 4E). We further performed correlation analysis using three custom gene panels composed of key regulators or markers for the maturation of mitochondria, sarcomeres, or cardiac excitability and calcium handling (Fig. 4F-H). We observed good correlation (r>0.6) between *Actn2* mutant and *Srf* KO experiments for each of these gene sets. These data suggest that *Actn2* regulates gene transcription in CM maturation by modulating SRF activity.

**Fig. 4.**
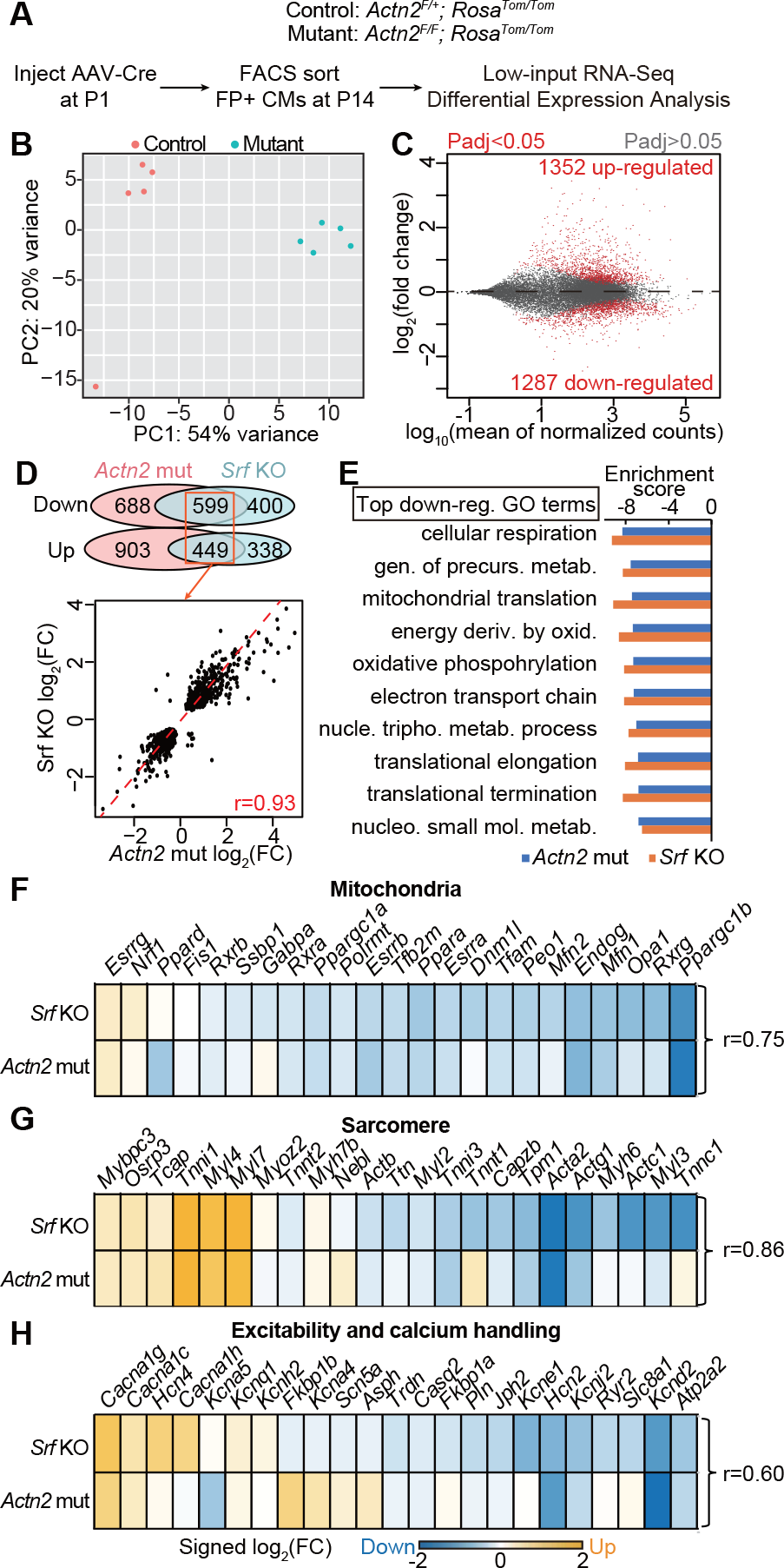
Cell-autonomous impact of *Actn2* on transcription. **(A)** Experimental design to evaluate cell autonomous regulation of the CM transcriptome downstream of *Actn2*. AAV-Cre was injected at low dose to P1 mice. Transduced CMs were isolated at P14, FACS-purified, and analyzed by RNA-seq. Control and mutant mice had the indicated genotypes. **(B)** PCA plot of RNA-seq results. **(C)** MA plot of RNA-seq results. **(D)** Venn diagram of overlapping differentially expressed genes in *Actn2* mutation and *Srf* KO experiments (top) and correlation analysis between fold changes of the shared differentially expressed genes (bottom). **(E)** Top gene ontology terms enriched among genes down-regulated in *Actn2* mutation and *Srf* KO experiments. Negative scores mean down-regulation. **(F-H)** Correlation analysis of key mitochondrial **(F)**, sarcomeric **(G),** and electrophysiological **(H)** genes between *Actn2* mutation and *Srf* KO experiments. r, Pearson correlation coefficient.

The polymeric state of the actin cytoskeleton regulates nuclear localization and transcriptional activity of SRF cofactors MRTFA/B. Thus, we tested the hypothesis that ACTN2 modulates actin polymerization and MRTF localization. We injected a high dose of AAV-Cre into *Actn2*^*F/+*^ and *Actn2*^*F/F*^ at P1 and collected heart tissue at P7 for biochemical analysis of actin. This approach circumvented perturbation of the actin cytoskeleton during CM isolation and FACS purification. Although *Actn2* mutation strongly reduced F-actin (Fig. 2A-B), total actin protein levels were not changed (Fig. 5A). These data suggested an increase in G-actin.

**Fig. 5.**
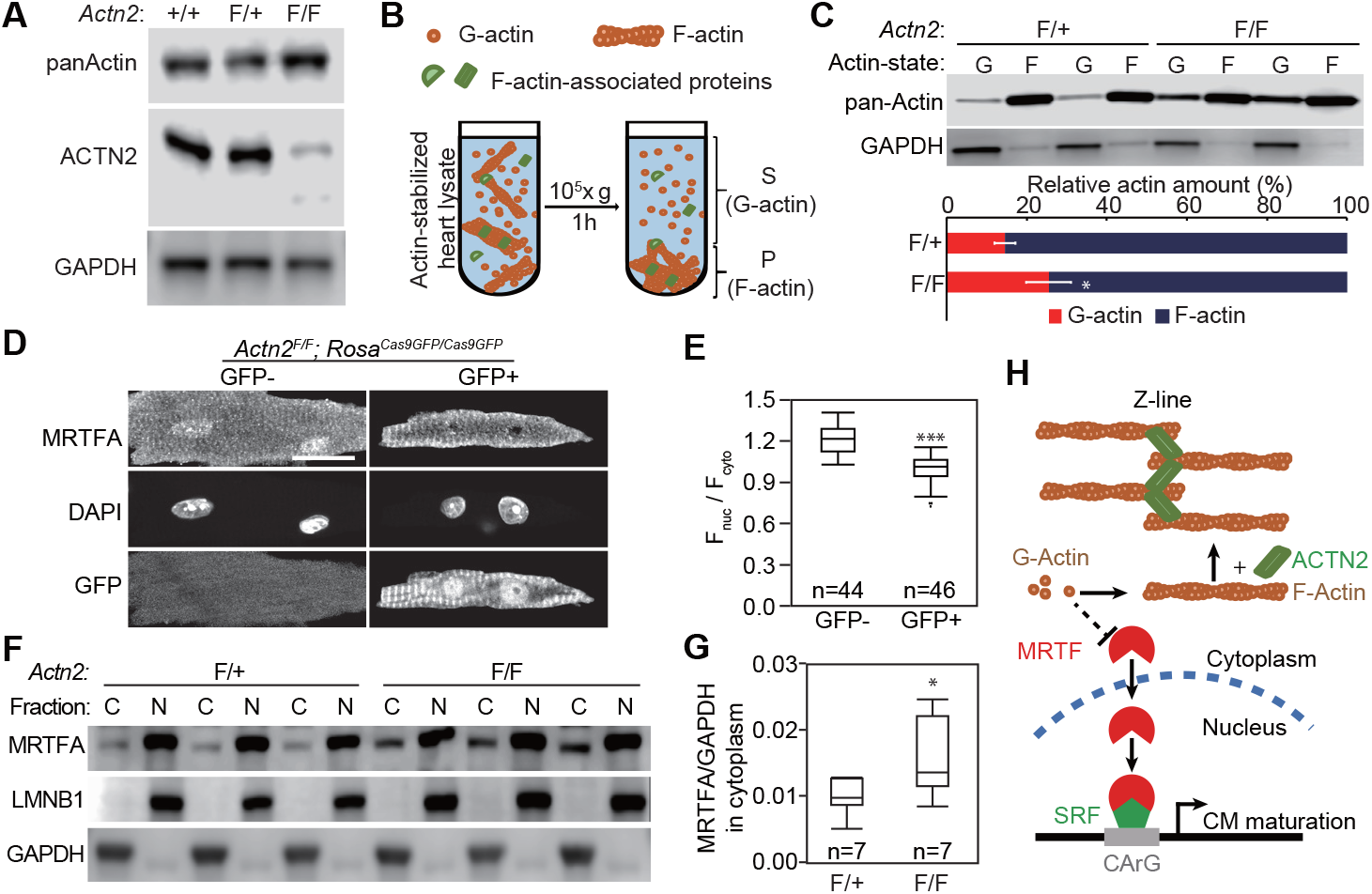
Impact of *Actn2* mutation on actin depolymerization and MRTF mislocalization. **(A)** Western blot analysis of total actin in P7 heart tissues treated with high dose AAV-Cre at P1. **(B)** Workflow of actin pelleting experiment. **(C)** Western blot (top) and densitometric quantification (bottom) of F-actin:G-actin ratio by actin pelleting analysis. Error bar is SD. n=4 hearts per group. High-dose AAV-Cre was injected at P1 and the heart tissues were collected at P7. **(D)** MRTFA localization in CMs isolated from hearts with mosaic *Actn2* mutation. Cells were immunostained with an MRTFA antibody and imaged using a confocal microscope. Bar = 20 μm. **(E)** Fluorescence intensity ratio quantification between nuclear and cytoplasmic MRTFA signals. **(F)** Western blot analysis of MRTFA, LMNB1 (nuclear marker), and GAPDH (cytoplasmic marker) in nuclear (N) and cytoplasmic (C) fractions. High-dose AAV-Cre was injected at P1 and the heart tissues were collected at P7. **(G)** Quantification of MRTFA western blot signal in the cytoplasmic fraction, normalized to GAPDH. **(H)** Model for ACTN2-MRTF-SRF sarcomere-to-nucleus signaling. ACTN2 promotes F-actin assembly into sarcomeres and decreases G-actin level, which allows MRTF to enter the nucleus and stimulate transcription through SRF. Box plots and statistical analyses are the same to **Fig. 2**.

To directly measure G-actin levels in heart tissues, we adopted an actin pelleting assay (*20*), in which ultracentrifugation pelleted F-actin while leaving G-actin in the supernatant (Fig. 5B). As a positive control, we treated WT heart lysates with Swinholide A, which severs F-actin and triggers its depolymerization (*21*). This treatment significantly increased the amount of actin in the supernatant (Fig. S5A-B), validating this assay for analysis of heart tissue. The actin pelleting assay showed that actin was predominantly assembled into F-actin in control heart tissue (Fig. 5C and Fig. S5C). In *Actn2* mutant heart tissue, G-actin was significantly more abundant (Fig. 5C).

We next assessed the impact of *Actn2* mutation on MRTF localization. Previous studies have reported conflicting results about MRTF localization in CMs (*22*, *23*), likely due to its low protein expression levels and limited signal-to-noise ratio of available antibodies in tissue sections. To circumvent these problems, we generated an AAV vector expressing MRTFA tagged with GFP. WT hearts transduced with AAV-MRTFA-GFP were perfused with Hoechst dye to label nuclei and then imaged *in situ* by confocal optical sectioning. This process avoided potential artifacts arising from tissue processing, CM isolation, or non-specific antibodies. This method clearly revealed that MRTFA is localized to CM nuclei at both P7 and P30 (Fig. S6A). To validate this finding for endogenous MRTFA protein, we stained wild-type dissociated CMs with a widely accepted antibody (*24*). We observed similar nuclear localization of endogenous MRTFA. CMs from the same hearts transduced by AAV-MRTFA-GFP showed dramatically increased nuclear staining compared to non-transduced CMs from the same heart (Fig. S6B), strongly suggesting that this immunolocalization is authentic.

Having established that MRTFA is normally localized to CM nuclei, we evaluated the effect of *Actn2* mutation and consequent elevation of G-actin on MRTFA localization. We treated *Actn2*^*F/F*^*;Rosa*^*Cas9GFP/Cas9GFP*^ mice with low dose AAV-Cre and immunostained CMs isolated from these hearts. Control and mutant CMs were stained in the same dish and imaged using the same confocal parameters on the same day to avoid immunostaining or imaging batch effects. Control and mutant CMs were distinguished by GFP signal. MRTFA nuclear and cytoplasmic signals were quantified by fluorescence intensity. We found that the nuclear to cytoplasmic MRTFA signal ratio was reduced in *Actn2* mutant CMs (Fig. 5D-E). To validate this result using an orthogonal method that circumvented potential confounding effects of CM isolation, we performed subcellular fractionation of *Actn2*^*F/F*^*;Rosa*^*Cas9GFP/Cas9GFP*^ and *Actn2*^*F/+*^*;Rosa*^*Cas9GFP/Cas9GFP*^ hearts treated with high dose AAV-Cre at P1 and collected at P7. Nuclear and cytoplasmic markers LMNB1 and GAPDH were found in the expected subcellular fractions (Fig. 5F), confirming clean separation of these compartments. MRTFA was enriched in the nuclear fraction. Cytoplasmic MRTFA was greater in *Actn2* mutant hearts compared to controls (Fig. 5F-G). The observed change in cytoplasmic MRTFA likely under-represents the actual change in Actn2 mutant cells because of the incomplete depletion of ACTN2^WT^, and the diluting effect of MRTFA in non-myocytes and non-transduced CMs.

Together, these data indicate that ACTN2 promotes actin polymerization, which reduces G-actin and enhances nuclear localization of MRTF in WT CMs (Fig. 5H). In contrast, *Actn2* mutation impairs actin polymerization and elevates G-actin, thereby reducing MRTF nuclear localization.

### Perturbation of CM maturation by MRTFA inhibition

To directly test the role of MRTF in CM maturation, we used AAV to express a dominant-negative mutant of MRTFA (dnMRTFA) (*25*). dnMRTFA lacks the transactivation domain (TAD, Fig. 6A) and competes with endogenous MRTFA and MRTFB for interaction with SRF. As a result, this approach inhibits both MRTF isoforms. Although dnMRTFA retains the G-actin-binding RPEL domains, which mediate G-actin regulation of MRTFA nuclear export, dnMRTFA-GFP was found mainly in the cell nucleus in all CMs that we examined by *in situ* imaging (Fig. 6B). This confirmed that the low G-actin level in CMs was insufficient to perturb nuclear dnMRTFA localization. We observed severe T-tubule defects in dnMRTFA-treated CMs (Fig. 6B-C). These cells also exhibited perturbed ACTN2 organization, characterized by the presence of longitudinal ACTN2 staining (Fig. 6D) and decreased distance between transverse ACTN2 patterns (Fig. 6E). Maturational CM hypertrophy was also perturbed by dnMRTFA (Fig. 6F). These morphological phenotypes were almost identical to those observed in *Srf* KO CMs (*2*), indicating a key role of MRTF in the activation of the essential SRF activity and the organization of ACTN2 for proper CM maturation.

**Fig. 6.**
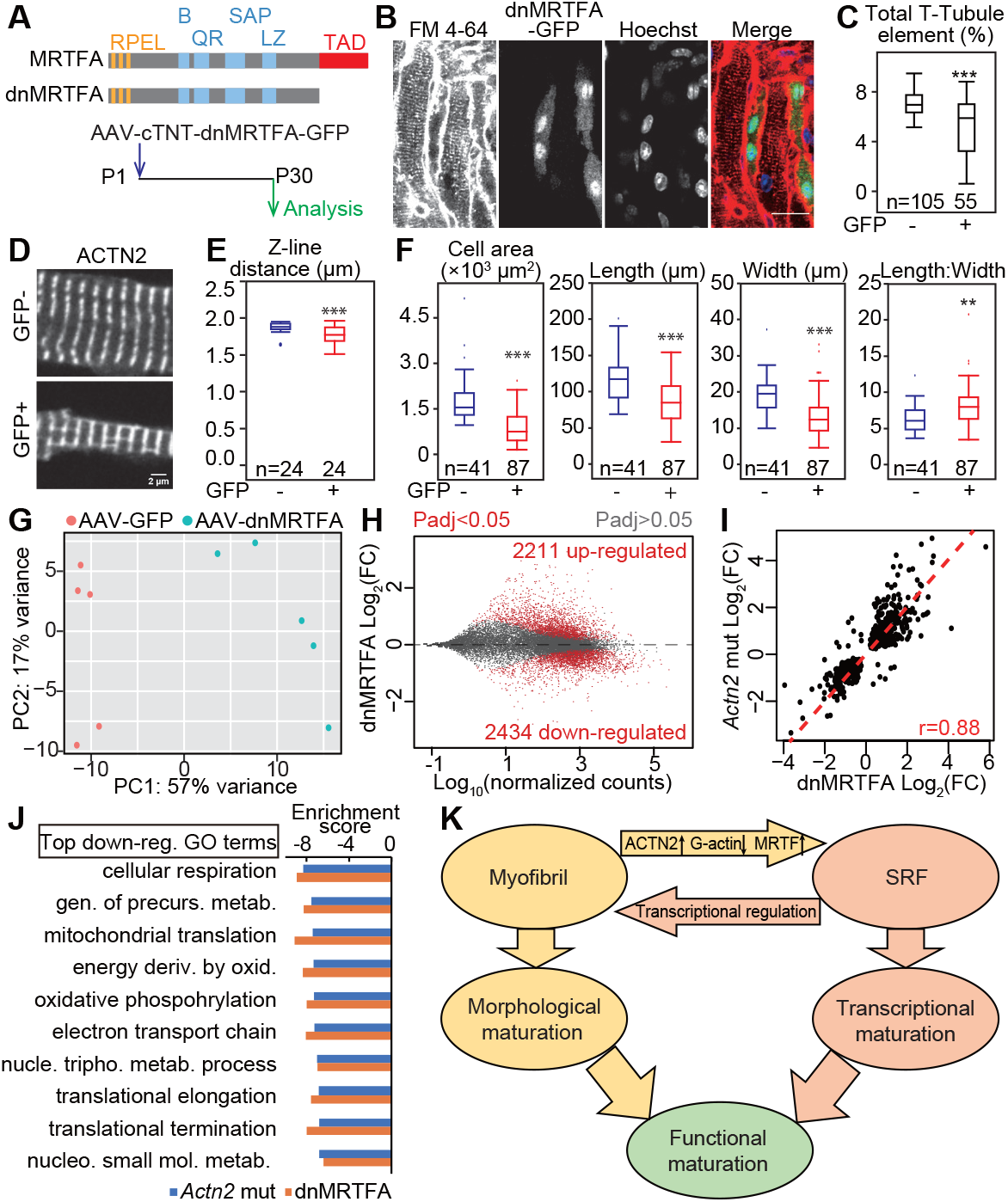
Perturbance of CM maturation by dominant-negative MRTFA . **(A)** Design of dnMRTFA-GFP overexpression experiments. RPEL, actin binding domain; TAD, transactivation domain. B, basic region. QR, glutamine-rich region. SAP, SAP domain. LZ, leucine zipper domain. **(B)** Confocal optical sectioning of hearts with mosaic dnMRTFA-GFP transduction. Hearts were perfused with Hoeschst dye and FM4-64. Bar = 20 μm. **(C)** AutoTT quantification of T-tubule content in dnMRTFA-treated hearts. **(D)** Immunofluorescent staining of ACTN2 in isolated CMs. **(E)** Quantification of Z-line distance in **(D)**. **(F)** Size and aspect ratio of dnMRTF-GFP and control CMs that were isolated from hearts with mosaic dnMRTFA expression. **(G-H)** Gene expression analysis of control and dnMRTFA CMs. Mice were treated with low dose AAV-dnMRTFA-GFP or AAV-GFP. GFP+ CMs were purified by FACS. RNA expression was measured by RNA-seq. Expression profiles of individual samples are displayed as a PCA plot in **(G)**. Differential expression analysis of RNA-seq data are shown in **(H)**. **(I)** Correlation analysis of shared differentially expressed genes in *Actn2* mutation and dnMRTFA treatment experiments. r, Pearson correlation coefficient. **(J)** Top 10 GO terms that are down-regulated in dnMRTFA-treated CMs were shared with the top GO terms from down-regulated genes in *Actn2* mutant CMs. **(K)** Summary of mutual potentiation between sarcomere assembly and MRTF-SRF signaling to coordinate CM maturation. Box plots and statistical analyses are the same as **Fig. 2.**

We next performed FACS-RNA-seq to characterize the impact of dnMRTFA on transcription, using AAV-GFP treated CMs as control. Five biological replicates of control vs. treated CMs were well separated by PCA (Fig. 6G; Suppl. Table 1). Statistical analysis revealed 2434 down-regulated genes and 2211 up-regulated genes (Fig. 6H; Suppl. Data 1). Gene fold changes were highly correlated between dnMRTFA and *Actn2* mutation RNA-seq datasets (Fig. 6I), and the same top-ranked GO terms were down-regulated as assessed by GSEA (Fig. 6J).

These data demonstrate similar impacts of *Actn2* mutation and MRTF inhibition on transcription in CM maturation and support the hypothesis that they function in the same signaling pathway.

## Discussion

Sarcomeres are components of a specialized cytoskeleton that mediates the heart’s contractile activity. However, less is known about potential non-contractile, signaling functions of sarcomeres. Sarcomere Z-lines have been proposed as signal transduction hubs (*4*), but the molecular mechanisms are ill-defined. Z-line signals were mainly investigated in the context of cardiac pathology using manipulations that impacted overall heart function, so that interpretation of the results of these studies is complicated by secondary effects of the cardiac stress response. Consequently, whether and how Z-lines regulate signaling in a physiological context has remained an open question. Here we combined AAV-mediated genetic mosaic analysis (*9*) with a newly generated floxed allele of *Actn2* to efficiently disassemble sarcomeres in a small fraction of CMs, while maintaining normal global cardiac function. This approach revealed a cell-autonomous role for ACTN2 in T-tubule morphogenesis, mitochondria organization, and physiological CM hypertrophy during CM maturation. These effects on CM structural maturation were coupled with a profound cell-autonomous impact of ACTN2 on CM gene transcription through retrograde sarcomere-to-nucleus signaling.

We identified at least one molecular mechanism that contributes to sarcomere-to-nucleus signaling: ACTN2 activates MRTF-SRF signaling by promoting actin polymerization, which is essential for CM maturation during heart development. Although the effects of actin dynamics on MRTF-SRF signaling are well-established in many cell types, it was unclear whether this mechanism exists in CMs due to the specialization of the CM actin cytoskeleton. In this study, we showed that actin polymerization into sarcomeres does indeed regulate MRTF nuclear localization and SRF activity. Under normal conditions, efficient sarcomere assembly in CMs results in a low G-actin concentration, nuclear MRTF enrichment, and activation of sarcomere gene expression through SRF. However, impairing sarcomere assembly by mutating *Actn2* elevates G-actin, reducing nuclear MRTF enrichment, and impeding CM maturation by disrupting MRTF-SRF signaling. Together with our recent discovery that SRF signaling activates sarcomere assembly (*2*), these data indicate mutual potentiation between sarcomere assembly and SRF signaling. This positive feedback loop ensures robust and coordinated maturation of both CM morphology and the CM transcriptional program (Fig. 6K).

*Actn2* mutation resulted in additional transcriptional dysregulation that was not observed in SRF-depleted CMs, suggesting that retrograde sarcomere-to-nucleus signaling could involve molecular pathways in addition to SRF-MRTF. Other ACTN2 binding partners may mediate these additional signaling pathways. For example, CSRP3 (also known as MLP) was reported to shuttle between Z-lines and nucleus (*26*), and CSRP3 knockout causes dilated cardiomyopathy in mice (*27*). Alternatively, impaired contractility in *Actn2* mutant CMs may induce these additional transcriptional changes. Future studies are warranted to fully define the mechanisms that govern ACTN2-dependent, retrograde sarcomere-to-nucleus signaling.

ACTN2-MRTF-SRF signaling may be relevant to the pathogenesis of human cardiomyopathies. Here, we interfered with sarcomere-to-nucleus signaling by mutating *Actn2*. We hypothesize that human sarcomere gene mutations that cause dilated cardiomyopathy or hypertrophic cardiomyopathy, including mutations in *ACTN2*, also impact this signaling pathway. While mechanistic studies of these sarcomere gene mutations have focused on their impact on contractile function or calcium binding, our study suggests that a subset of these mutations could cause disease by perturbing sarcomere-to-nucleus signaling. This hypothesized mechanism also predicts that disruption of the coordinated CM maturation program contributes to the pathogenesis of these genetic cardiomyopathies.

CMs that are derived from non-myocytes or stem cells are widely used in disease modeling, pharmacological tests, tissue engineering, and cell therapy trials. These cells exhibit immature phenotypes *in vitro* and do not fully recapitulate native CMs, imposing a major bottleneck on cardiac regenerative medicine (*1*). Many approaches have been developed to promote CM maturation, and an improved ACTN2 staining pattern has been treated as a key hallmark of ultrastructural maturation. Here we show that in addition to its structural role ACTN2 regulates signal transduction, providing a novel insight behind the effort to improve ACTN2 organization. Thus, we propose that sarcomere maturation is a key prerequisite to promote overall CM maturation.

## Materials and Methods

### Mouse strains

All animal strains and procedures were approved by the Institutional Animal Care and Use Committee of Boston Children’s Hospital. *Rosa*^*Cas9GFP/Cas9GFP*^ (Stock No: 026175) (*14*) and *Rosa*^*Tomato/Tomato*^ (Stock No: 007914) (*16*) were imported from the Jackson Laboratory. *Actn2*^*F*^ mice were generated by homologous recombination in ES cells as shown in Figure 1A. All mice that were used in this study were on a mixed genetic background.

### Plasmids

AAV-cTNT-Cre, AAV-cTNT-GFP and AAV-cTNT-GFP-version2 were described previously (*2*). Wild-type *Actn2* and mutant *Actn2*^*ΔE2-6*^ coding sequences were synthesized (Integrated DNA technologies, gBlock) and cloned into AAV-cTNT-GFP-version2 to generate AAV-ACTN2^WT^-GFP and AAV-ACTN2^ΔE2-6^-GFP plasmids. Murine MRTFA coding sequence was acquired from GE Healthcare Dharmacon (MMM1013-202859509). Full-length *Mrtfa* and dn*Mrtfa* sequences were PCR amplified using this template and cloned into AAV-cTNT-GFP-version2 to generate AAV-MRTFA-GFP and AAV-dnMRTFA-GFP plasmids.

### AAV production and injection

AAV9 was prepared as previously described (*2*, *8*) with modifications. In brief, 140 μg AAV-ITR, 140 μg AAV9-Rep/Cap, and 320 μg pHelper (pAd-deltaF6, Penn Vector Core) plasmids were produced by maxiprep (Invitrogen, K210017) or Gigaprep (Invitrogen, K210009XP) and triple transfected into 10 15-cm plates of HEK293T cells using PEI transfection reagent (Polysciences, 23966- 2). 60 h after transfection, cells were scraped off of plates, resuspended in lysis buffer (20 mM Tris pH 8, 150 mM NaCl, 1 mM MgCl2, 50 μg/ml Benzonase) and lysed by three freeze-thaw cycles. AAV in cell culture medium was precipitated by PEG8000 (VWR, 97061-100), resuspended in lysis buffer and pooled with cell lysates. AAV was purified in a density gradient (Cosmo Bio USA, AXS-1114542) by ultracentrifugation (Beckman, XL-90) with a VTi-50 rotor and concentrated in PBS with 0.001% pluronic F68 (Invitrogen, 24040032) using a 100 kD filter tube (Fisher Scientific, UFC910024). AAV titer was quantified by real-time PCR using a fragment of the TNT promoter DNA to make a standard curve. PCR primers for AAV quantification were 5’-TCGGGATAAAAGCAGTCTGG-3’ and 5’-CCCAAGCTATTGTGTGGCCT-3’. SYBR Green master mix (Invitrogen, 4368577) was used in this quantification.

AAV was injected into P1 pups subcutaneously. The P1 pups were anesthetized in an isoflurane chamber before injection. AAV dosage was normalized based on body weight and was described in Figure 2A. 3.6×10^8^ viral genome per gram body weight (vg/g) was used in all mosaic analyses.

### Echocardiography

Echocardiography was performed on a VisualSonics Vevo 2100 machine with Vevostrain software by an investigator blinded to genotype or treatment group . Animals were awake during this procedure and held in a standard handgrip.

### Histological analysis

After the animals were euthanized by CO_2_, hearts were harvested immediately and fixed by 4% paraformaldehyde overnight at 4 °C. Fixed hearts were cryoprotected by soaking in 15% sucrose followed by 30% sucrose at 4 °C. Hearts were embedded in tissue freezing medium (General Data, TFM-5). 10 μm cryo-sections were cut using a cryostat (Thermo Scientific, Microm HM 550).

Fast green and sirius red staining was performed on frozen sections. Sections were first washed with PBS for 5min, fixed with pre-warmed Bouin’s solution (Sigma, HT10132) at 55 °C for 1h and washed in running water. The sections were then stained with 0.1% Fast green (Millipore, 1040220025) for 10 minutes, washed with 1% acetic acid for 2 minutes, and rinsed with running water for 1 minutes. The sections were then stained with 0.1% sirius red (Sigma, 365548) for 30 minutes and washed with running water for 1 minute. The slides were treated with 95% ethanol once for 5min, twice with 100% ethanol for 5min, and twice in Xylene for 5min before being mounted with Permount (Fisher Scientific, SP15-500). Bright-field images of stained tissue sections were taken with a BZ-X700 all-in-one microscope (Keyence).

### In situ confocal imaging

In situ T-tubule labeling was performed as previously described (*15*). Hearts were dissected from euthanized animals and cannulated on a Langendorff apparatus. Hearts were perfused withperfusion buffer [10 mM Hepes (pH 7.4), 120.4 mM NaCl, 14.7 mM KCl, 0.6 mM KH_2_PO_4_, 0.6 mM Na_2_HPO_4_, 1.2 mM MgSO_4_, 4.6 mM NaHCO_3_, 30 mM Taurine, 10 mM 2,3-butanedione monoxime, 5.5 mM glucose] at room temperature for 10 min. Where indicated, FM 4-64 (Invitrogen, 13320), TMRM (source), or Hoechst 33342 (Invitrogen) dyes were added to the perfusion buffer at 2 μg/ml, 2 nM, and 4 μg/ml, respectively.

After in situ labeling, the heart was removed from the perfusion system, positioned on a glass-bottom dish, and immediately imaged using an inverted confocal microscope.

### Cardiomyocyte isolation

Cardiomyocytes were isolated by retrograde collagenase perfusion using an established protocol (*28*). In brief, heparin-injected mice were anesthetized in an isoflurane chamber. Hearts were isolated and cannulated onto a Langendorff perfusion apparatus. 37 °C perfusion buffer was first pumped into the heart to flush out blood and equilibrate the heart. Collagenase II (Worthington, LS004177) was next perfused into the heart for 10 min at 37 °C to dissociate cardiomyocytes. The apex was cut from the digested heart, gently dissociated into single cardiomyocytes in 10% FBS/perfusion buffer, and filtered through a 100 μm cell strainer to remove undigested tissues.

### Immunofluorescence

To prepare cells for immunofluorescence, isolated cardiomyocytes were concentrated by 20 × g centrifugation for 5 min and re-suspended in cell culture medium [DMEM (Gibco), 10% FBS, pen/strep (Gibco), 10 μM Blebbistatin]. Cardiomyocytes were cultured on laminin-coated coverslips for ~1h at 37 °C with 5% CO^2^ to allow cells to attach to the coverslips.

Next, immunofluorescent labeling was performed as described (*8*, *11*, *29*, *30*). Cardiomyocytes were fixed with 4% paraformaldehyde for 10~20 min, permeabilized by 0.1% Triton-100/PBS for 10 min, and blocked in 4% BSA/PBS (blocking buffer) at 4°C overnight. Then the cells were incubated with primary antibodies diluted in blocking buffer overnight at 4 °C. After washes with blocking buffer, the cells were incubated with secondary antibodies and dyes at room temperature for 2 h. The cells were next washed with PBS and mounted with ProLong Diamond antifade mountant (Invitrogen, 36961) before imaging. All antibodies and dyes are listed in Supplementary Table 2.

### Fluorescence imaging and analysis

Confocal fluorescent images were taken using Olympus FV3000RS inverted laser scanning confocal microscope with a 60X/1.3 silicone-oil objective. Fluorescence intensity was measured using ImageJ. AutoTT (*18*) was used to quantify T-tubule and sarcomere organization. Cell size and shape were manually measured on maximally projected images.

### Contractility measurement

Calcium was re-introduced into isolated cardiomyocytes by treating cells with a series of 5 ml perfusion buffers containing 60 nM, 400 nM, 900 nM, and 1.2 μM CaCl_2_. At each step, cardiomyocytes were allowed to sediment for 5 min at room temperature before cells being transferred to the next buffer with higher calcium concentration.

For the contractility assay, cardiomyocytes were plated on laminin-coated coverslips in MEM media with 10 mM 2,3-butanedione monoxime and incubated for 30 min at 37 °C and 5% CO_2_. The coverslips were then transferred into a 37 °C flow chamber with 1.2 μM CaCl_2_ perfusion buffer without BDM. FP- and FP+ cells were identified by epifluorescence imaging. Cardiomyocytes were electrically stimulated at 1 Hz and cell contraction was measured using an IonOptix system and a 40X objective. Contractile and non-contractile CMs were identified visually.

### Electron microscopy analysis after fluorescence-activated cell sorting (FACS-EM)

Isolated cardiomyocytes in suspension were fixed with 4% paraformaldehyde for 30 min at room temperature. The fixed cells were next filtered by passing through a 100 μm cell strainer, pelleted by centrifugation at 20 × g for 5 min at room temperature and resuspended in ~1 ml perfusion buffer. FACS was performed using a BD AriaII SORP cell sorter with a 100 μm nozzle. After FACS, the cells were fixed again in a mixture of 2% formaldehyde and 2.5% glutaraldehyde in 0.1 M Sodium Cacodylate buffer, pH 7.4 overnight at 4°C. The cell pellets were next processed through a routine TEM protocol at the Harvard Medical School EM core. Images were taken using a JEOL 1200EX - 80kV electron microscope. Because of the size and stiffness of cardiomyocytes, fixed adult cardiomyocytes clog the FACS machine. As a result, FACS-EM currently only works for CMs from P30 or younger mice.

### Real-time quantitative PCR (RT-qPCR) and reverse-transcription PCR (RT-PCR)

For RT-qPCR analysis using heart tissue, total RNA was purified using PureLink RNA Mini kit (Ambion, 12183025). Genomic DNA removal and reverse transcription was performed using the QuantiTech reverse transcription kit (Qiagen, 205311). Real-time PCR was performed using an ABI 7500 thermocycler.

For RT-qPCR using FACS sorted cells, isolated cardiomyocytes were filtered with a 100 μm cell strainer, pelleted by centrifugation at 20 × g for 5 min and resuspended in ~1 ml cold perfusion buffer. FACS was performed using a BD AriaII SORP cell sorter with a 100 μm nozzle and a sample collection cooling device. Immediately after FACS, cells were centrifuged at 13,000 rpm at 4°C to remove supernatant. Total RNA was purified using PureLink RNA Micro kit (Thermo Fisher, 12183016) and genomic DNA was removed by on-column DNase I digestion. Reverse transcription was performed using the SMART-Seq v4 Ultra Low Input RNA Kit (Clontech), which pre-amplified full-length cDNA before qPCR. Real-time PCR was performed using an ABI 7500 thermocycler.

RT-qPCR was performed using either SYBR Green or Taqman probes. For SYBR detection, *Nppa* detection primers were 5’-TTCCTCGTCTTGGCCTTTTG-3’ and 5’-CCTCATCTTCTACCGGCATC-3’. *Nppb* detection primers were 5’-GTCCAGCAGAGACCTCAAAA-3’ and 5’-AGGCAGAGTCAGAAACTGGA-3’. *Gapdh* (internal control) detection primers were 5’-AGGTCGGTGTGAACGGATTTG-3’ and 5’-TGTAGACCATGTAGTTGAGGTCA-3’. For Taqman quantification, *Actn2 and Gapdh* were measured using Mm00473657_m1 (Invitrogen, 4453320) and mouse GAPD endogenous control (Invitrogen, 4352339E) assays, respectively.

RT-PCR detection of *Actn2* splicing variants was performed using GoTaq PCR master mix (Promega) and primers: 5’-CCATCCATCCATCCATTTCTC-3’ and 5’-TCTTCCATCAGCCTCTCATTC-3’.

### Western blot analysis

Heart tissues were homogenized in RIPA buffer (25 mM Tris pH7.4, 150 mM NaCl, 1% Triton X-100, 0.5% Na Deoxycholate, 0.1% SDS) buffer supplemented with protease inhibitor cocktail and Micrococcal Nuclease at ~33.3 mg/ml concentration. Heart lysates were denatured in 2X SDS sample buffer at 70 °C for 10 min, separated on a 4%-12% gradient gel (Invitrogen, Bolt gels, NW04122BOX), transferred to a PDVF membrane, and blocked by 4% milk/TBST. Primary antibodies were incubated with the membrane overnight at 4°C, followed by four 15 min TBST washes. HRP-conjugated secondary antibodies were probed for 1~2h at RT, followed by four 15 min TBST washes. After adding Immobilon Western chemiluminescent HRP substrate (Millipore, WBKLS0500), chemiluminescence was detected using a Li-Cor C-DiGit blot scanner. Antibodies used in this study are listed in Supplementary Table 1.

### RNA-seq and data analysis

FACS-sorted CMs were centrifuged at 13000 rpm for 1 min and supernatant fluids were removed. Total RNA was extracted using PureLink RNA micro kit (Thermo Fisher, 12183016) with genome DNA removed through on-column DNase I digestion. 10 ng total RNA was reverse transcribed and full-length cDNA was specifically amplified by 8 PCR cycles using SMART-Seq v4 Ultra Low Input RNA Kit (Clontech) (*31*). RNA-seq libraries were constructed using Illumina’s Nextera XT kit and single-end sequence reads were obtained using an Illumina NextSeq 550 sequencer.

RNA-seq reads were aligned to mm10 by STAR (*32*) and reads counts were calculated by FeatureCounts (*33*). DESeq2 was used to perform statistical analysis of differential gene expression (*34*). An adjusted P value of 0.05 was used as the cutoff to identify differentially regulated genes. GO term analysis was performed using GSEA analysis with ranked gene lists (*35*). To visualize reads distribution on genomic loci, deeptools (*36*) were used to generate .bigwig files with normalized reads count and loaded to integrative genomics viewer (*37*).

### Actin pelleting analysis

Actin pelleting experiments were performed using the G-Actin/F-actin In Vivo Assay Biochem Kit (Cytoskeleton, Inc.) with modifications. In brief, heart tissues were homogenized in tissue lysis and actin stabilizing buffer (50 mM PIPES pH 6.9, 50 mM KCl, 5 mM MgCl^2^, 5 mM EGTA, 5% glycerol, 0.1% NP40, 0.1% Triton X-100, 0.1% Tween 20, 0.1% beta-mercaptoethanol) supplemented with ATP and protease inhibitors at 33 mg/ml concentration. The heart lysate was next incubated at 37 °C for 1 hr and centrifuged with a SW55Ti rotor for 1 hr at 33000 rpm at 20 °C. The pellet was dissolved in 8M Urea. The supernatant and pellets were diluted to the same volume in SDS sample buffer before Western Blot analysis.

As a positive control for actin pelleting, 1 μM Swinholide A was added into heart lysates before 37 °C incubation. Vehicle (ethanol) treatment was used as a control for Swinholide A treatment.

### Subcellular fractionation of heart tissue

Cytoplasmic and nuclear fractions of heart tissues were generated using the Subcellular Protein Fractionation Kit for Tissues (Life Technologies # 87790) with modifications. In brief, heart tissues were homogenized in 350 μl cytoplasm extract buffer, and spun at 3000g at 4 °C for 5min. The supernatant was treated as the cytoplasmic fraction. The pellet was resuspended and partially dissolved in 230 μl membrane extract buffer at 4 °C for 5 min. The lysate was next centrifuged at 3000g for 5 min. The pellet was resuspended in 90 μl nucleus extract buffer with micrococcal nuclease and dissolved at room temperature for 20 min. This mixture was next spun at 12000g for 5 min. The resulting supernatant was the nuclear fraction.

### Statistical Analysis

Statistical analysis and plotting were performed using JMP® software (SAS Institute). Student’s t-test was used to compare two groups. *P<0.05, **P<0.01, ***P<0.001. Numbers in parentheses in figures indicate non-significant P-values. RNA-seq differential expression analysis was performed using DESeq2 (34).

In box plots, the horizontal line within the box represents the median; the ends of the box represent the 25th and 75th quantiles; whiskers extend 1.5 times the interquartile range from the 25th and 75th percentiles; dots represent possible outliers. Bar chars were plotted as mean ± standard deviation.

## General

We appreciate the technical support by Dana-Farber Flow Cytometry Core, HMS Electron Microscopy Core, HMS Biopolymers Facility, and BCH Animal Facility. We thank Dr. Guido Posern for sharing MRTF antibody, Xiaoran Zhang for advice on bioinformatic analysis, Yulan Ai for technical assistance, and Justin King for manuscript editing. We thank all members in the labs of William Pu, Da-Zhi Wang and Sean Li for general discussion about this project.

## Author contributions

Y.G. conceived and designed the study. Y.G. and B.D.J. performed all experiments. B.M., E.C.T., and A.H.B generated and validated *Actn2*^*F*^ mice. Y.G., I.S. and G-C.Y. performed bioinformatic analysis. M.A.T and E.M.S provided critical support for MRTF experiments. Q.M. performed and analyzed echocardiograms. W.T.P. provided funding and overall supervision of this project. Y.G. and W.T.P wrote the manuscript.

## Data and materials availability

RNA-seq data have been deposited in the Gene Expression Omnibus (GEO) database under the accession codes GSE136096. All other data supporting the findings of this study are available within the article and its Supplementary information files or from the corresponding author upon reasonable request. *Actn2*^*F*^ mice were produced in Dr. Alan Beggs’s lab and could be obtained through a material transfer agreement.

---

**Fig. S1.**
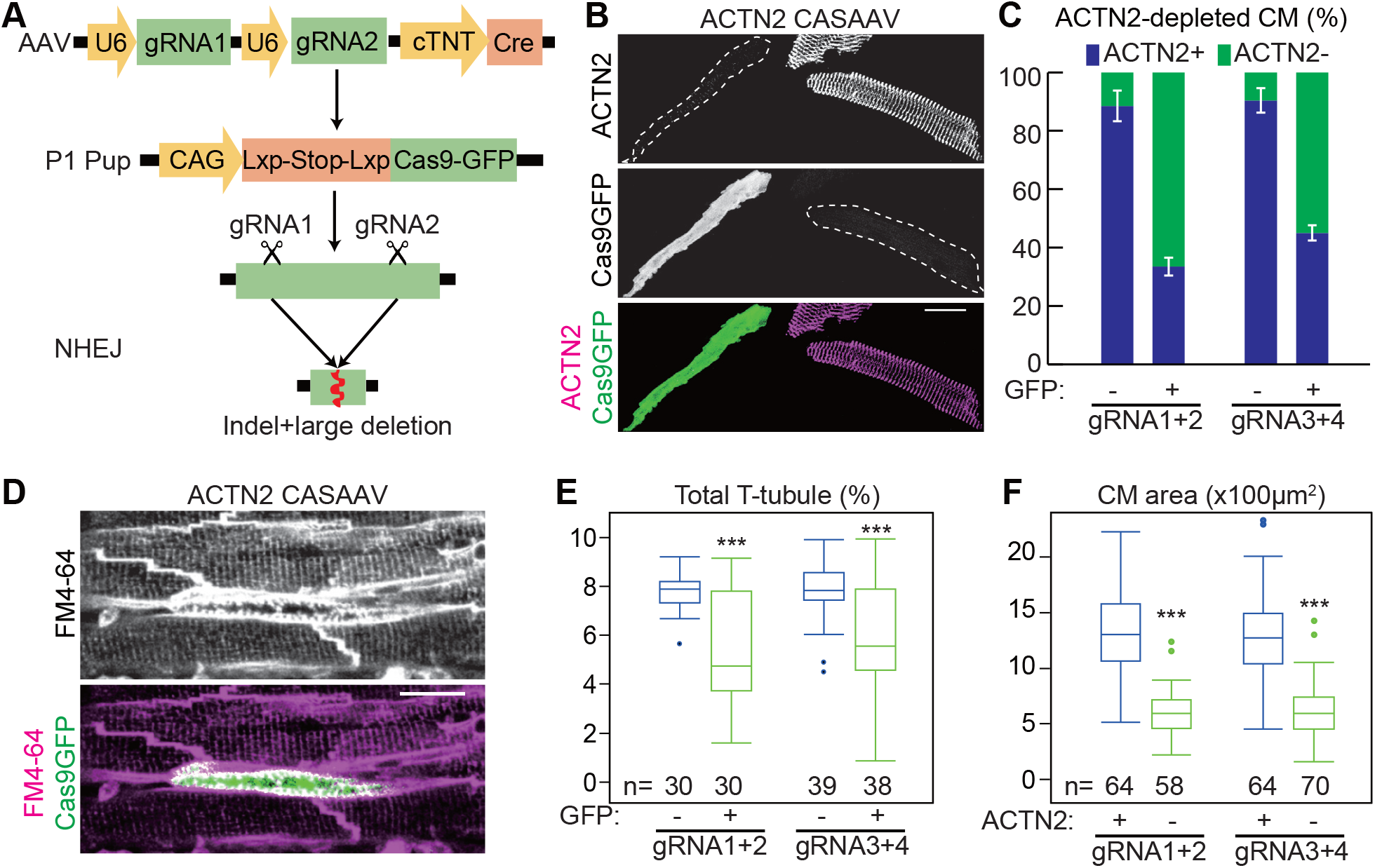
CASAAV-based mutagenesis of *Actn2* causes morphological defects in CM maturation. **(A)** CASAAV-mediated somatic mutagenesis. AAV9 drives expression of two gene-targeting guide RNAs and cardiomyocyte-specific expression of Cre. Cre activates expression of Cas9 and GFP from the Rosa26 locus. The gRNAs direct Cas9 to cleave the target gene at one or both gRNA-targeted sites, and introduce loss-of-function mutations. **(B-C)** Efficiency of ACTN2 depletion by CASAAV. AAV was injected at P1 and CMs were dissociated at P30 and immunostained for ACTN2 **(B)**. Cell boundaries are delin-eated by white dashed lines. ACTN2 depletion in CASAAV-treated CMs (GFP+) was quantified **(C)**. n=3 hearts per group; error bar, SD. **(D-E)** Effect of ACTN2 depletion on T-tubule morphology was analyzed by *in situ* imaging. *Actn2* CASAAV-treated P30 hearts were perfused with FM4-64, which labels the sarcolemma, and optically sectioned by confocal microscopy **(D)**. T-tubule content was quantified using AutoTT **(E)**. **(F)** Effect of ACTN2 depletion on CM size. Projected area of P30 dissociated CMs was plotted for cells with and without ACTN2 immunoreactivity. Student’s *t*-test:***p<0.001. Scale bars in **(B)** and **(D)** are 20 μm.

**Fig. S2.**
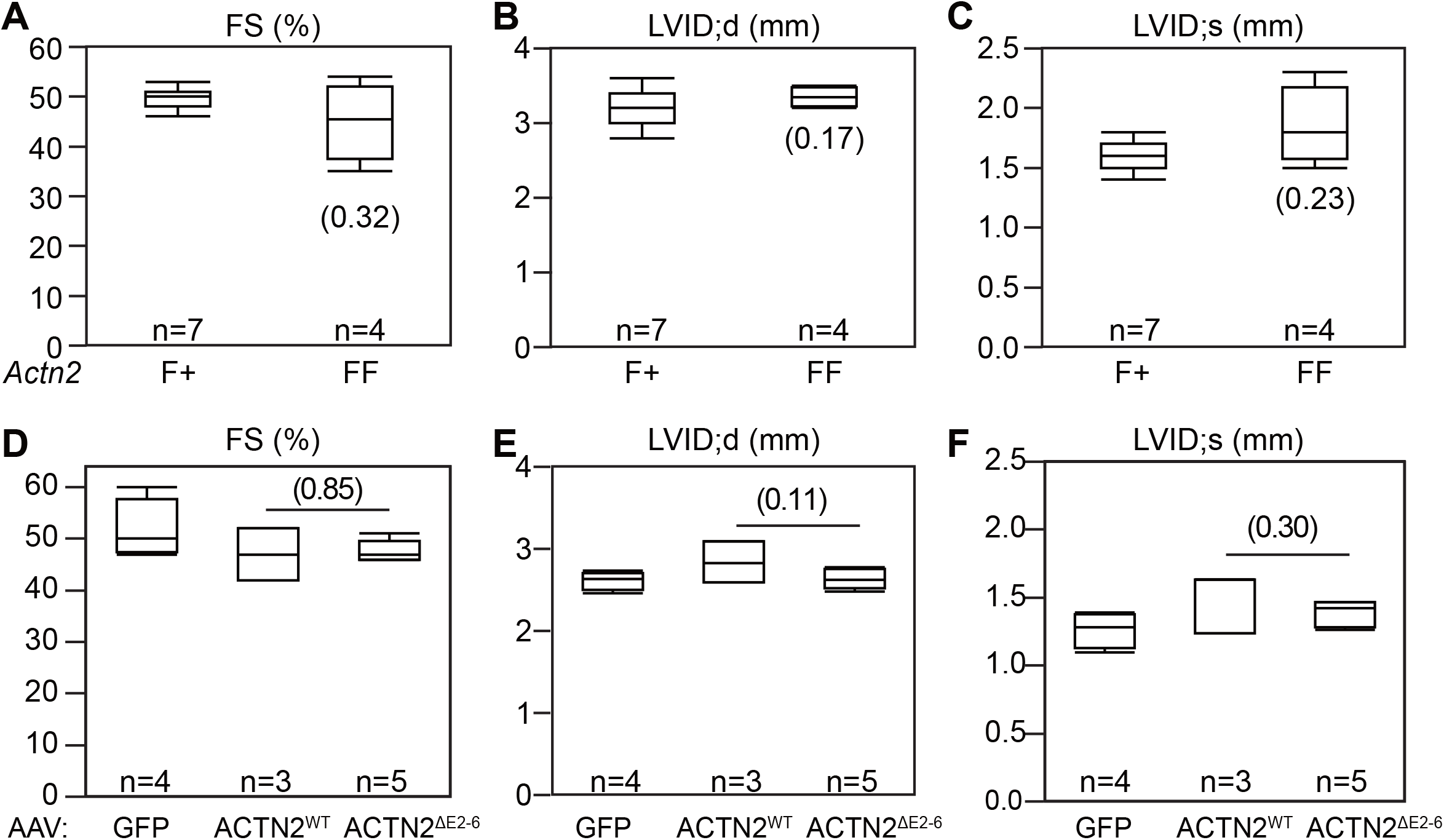
Echocardiographic measurement of heart function. **(A-C)** Analysis of 4-6 month old mice with no Cre treatment. **(D-F)** Analysis of P30 mice treated at P1 with high-dose AAVs expressing GFP, ACTN2^WT^-GFP, or ACTN2^ΔE2-6^-GFP. Student’s *t*-test: Non-significant P values in parenthesis.

**Fig. S3.**
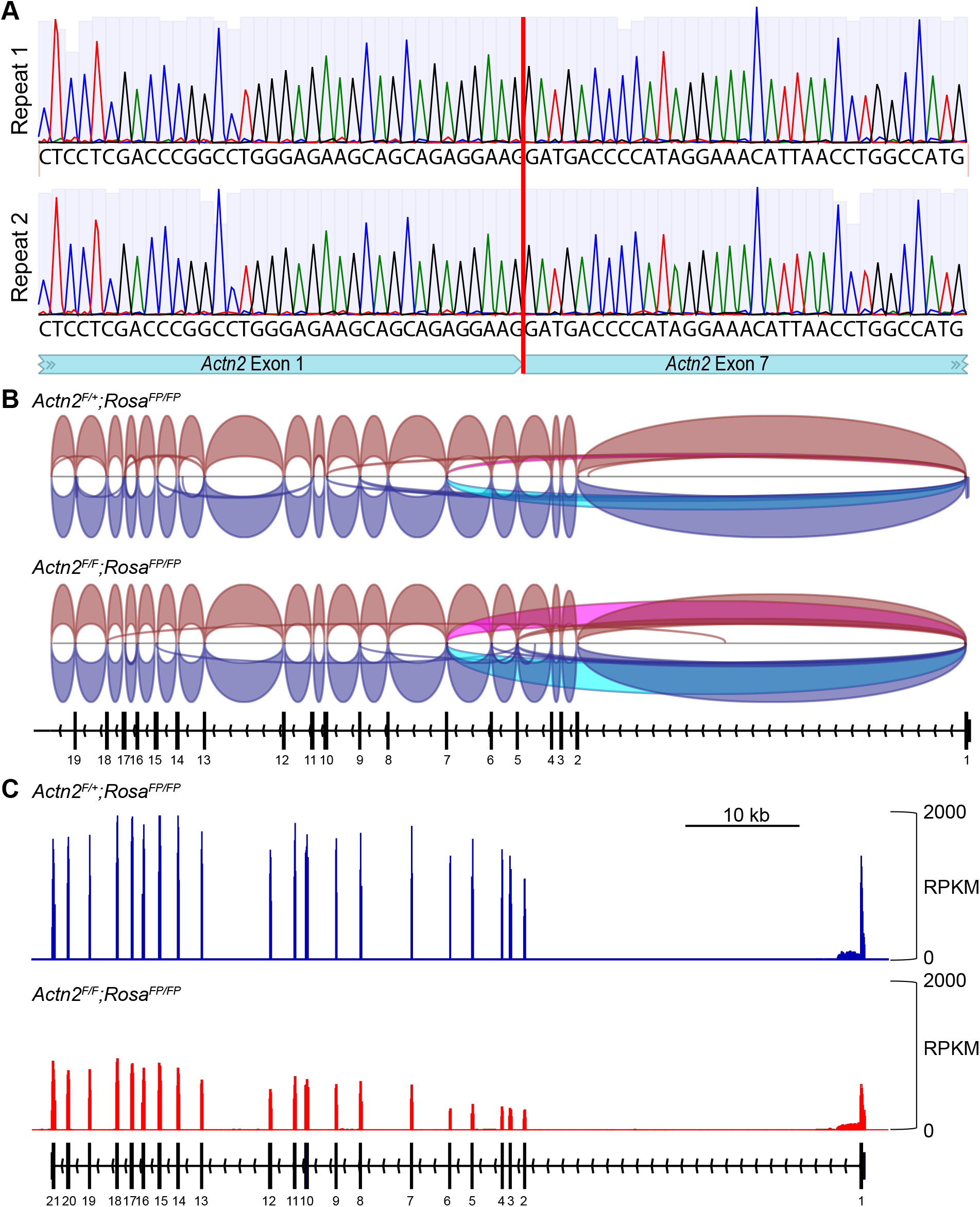
Sequencing analysis of *Actn2* transcripts. **(A)** Sanger sequencing of RT-PCR products showing ectopic splicing junction between *Actn2* exon 1 and 7, viewed using Benchling. **(B)** Profiling *Actn2* splicing junctions by RNA-seq using integrative genome viewer. Each splice junction is represented by an arc from the beginning to the end of the junction. The thickness of the arc represents the reads counts. Junctions from the plus strand are colored red and extend above the center counts. Junctions from the plus strand are colored red and extend above the center line. Junctions from the minus strand are blue and extend below the center line. The ectopic exon1-7 junction is highlighted in magenta and cyan. **(C)** The distrubtion of RNA-seq reads aligned to *Actn2* with a bin size of 50bp. The heights of the bars represent normalized reads count.

**Fig. S4.**
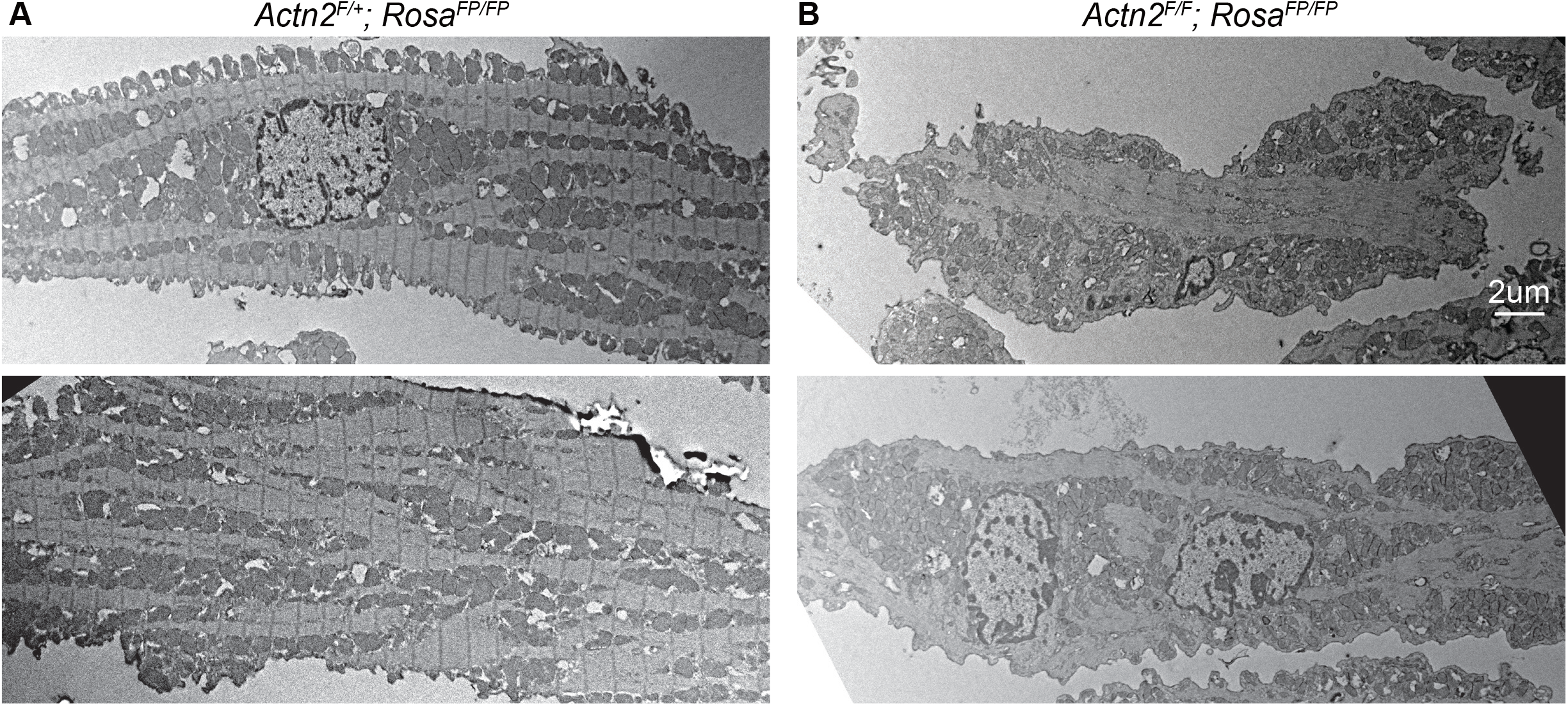
Electron microscopy analysis. *Actn2*^*F/+*^*; Rosa*^*FP/FP*^ **(A)** and *Actn2*^*F/F*^*; Rosa*^*FP/FP*^ **(B)** mice were treated with AAV-Cre at P1. At P30, CMs were isolated from these mice, fixed by PFA, and subjected to FACS purification of FP+ cells. FP+ Control **(A)** and mutant CMs **(B)** were next processed and imaged by electron microscopy.

**Fig. S5.**
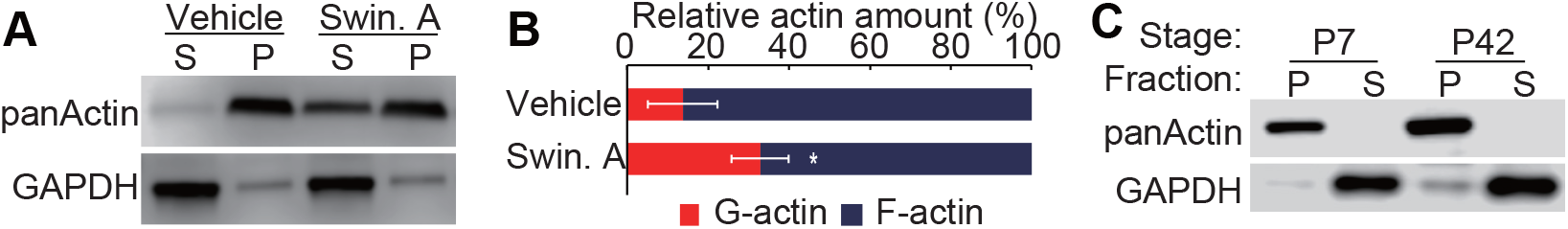
Assessment of actin polymerization by the actin pelleting assay. **(A)** Western blot showing increased actin in supernatant (S) com-pared to pellet (P) upon 5 μM swinholide A treatment of WT heart lysate. **(B)** Densitomectric quantification of relative G-actin (supernatant) and F-actin (pellet) in WT heart lysates by actin pelleting. n=3; error bar, SD. Student’s *t*-test: *, p<0.05. **(C)** Actin pelleting analysis of P7 and P42 WT heart lysates.

**Fig. S6.**
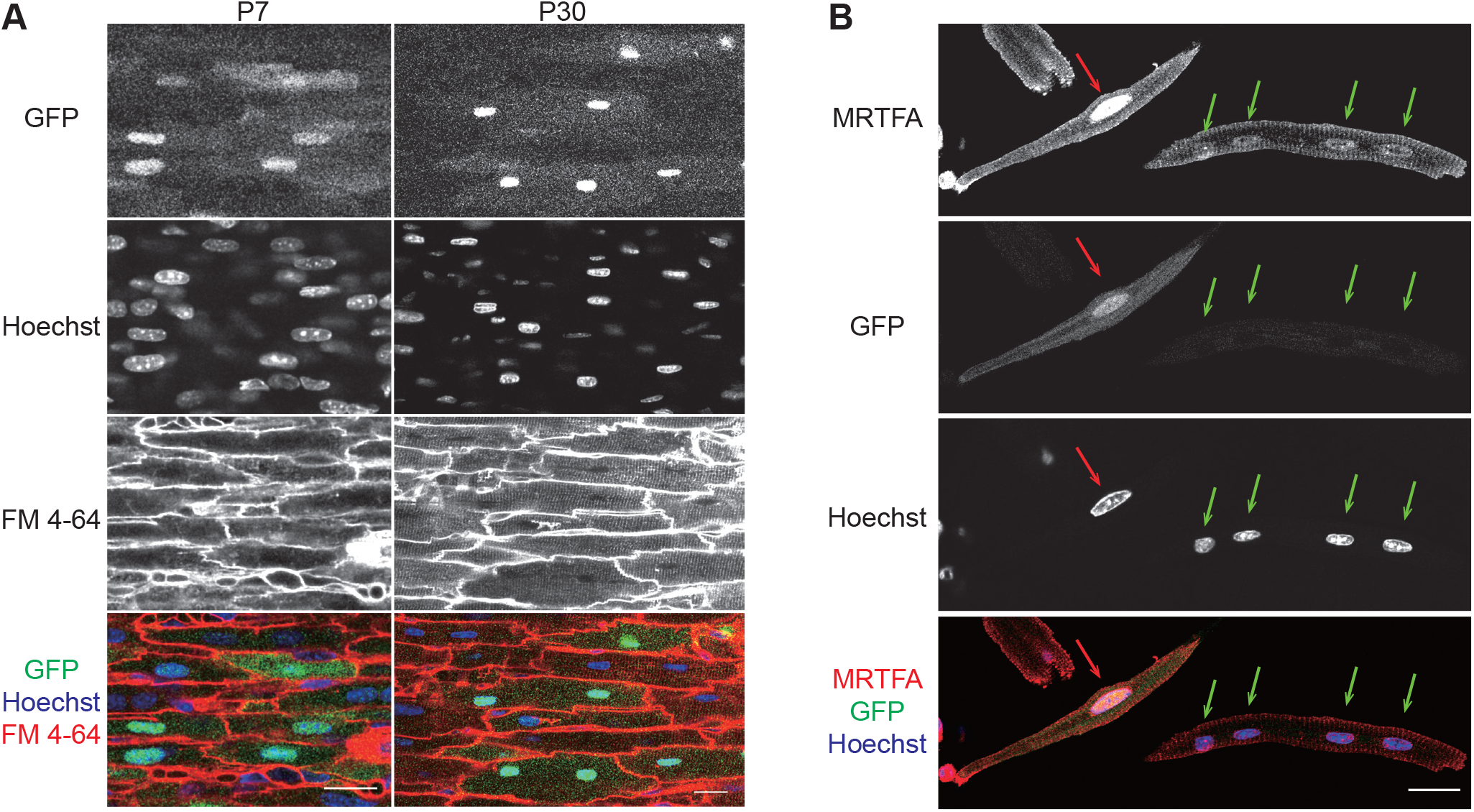
MRTFA localized to the nucleus of normal CMs. **(A)** In situ imaging MRTFA-GFP localization. Wild-type hearts transduced with AAV-MRTFA-GFP were perfused with Hoeschst and FM 4-64 dyes, which label nuclei and sarcolemma, respectively. The hearts were then optically sectioned using a confocal microscope. **(B).** Immunolocalization of endogenous and overexpressed MRTFA. AAV-MRTFA-GFP transduced hearts were dissociated and CMs were immunostained with MRTFA antibody. Untransduced CMs (GFP-) showed weak MRTFA immunoreactivity in their nuclei (green arrows). Transduced CMs (GFP+) showed stronger MRTFA predominantly in the nucleus. (red arrow). Scale bar, 20 μm.

## References

1. X. Yang, L. Pabon, C. E. Murry, Engineering Adolescence. Circ. Res. 114, 511–523 (2014).

2. Y. Guo, B. D. Jardin, P. Zhou, I. Sethi, B. N. Akerberg, C. N. Toepfer, Y. Ai, Y. Li, Q. Ma, S. Guatimosim, Y. Hu, G. Varuzhanyan, N. J. VanDusen, D. Zhang, D. C. Chan, G.-C. Yuan, C. E. Seidman, J. G. Seidman, W. T. Pu, Hierarchical and stage-specific regulation of murine cardiomyocyte maturation by serum response factor. Nat. Commun. 9, 3837 (2018).

3. W. G. Pyle, R. J. Solaro, At the crossroads of myocardial signaling: the role of Z-discs in intracellular signaling and cardiac function. Circ. Res. 94, 296–305 (2004).

4. D. Frank, N. Frey, Cardiac Z-disc signaling network. J. Biol. Chem. 286, 9897–9904 (2011).

5. M. K. Vartiainen, S. Guettler, B. Larijani, R. Treisman, Nuclear actin regulates dynamic subcellular localization and activity of the SRF cofactor MAL. Science. 316, 1749–1752 (2007).

6. T. Kobayashi, L. Jin, P. P. de Tombe, Cardiac thin filament regulation. Pflugers Arch. 457, 37–46 (2008).

7. M. H. Mokalled, K. J. Carroll, B. K. Cenik, B. Chen, N. Liu, E. N. Olson, R. Bassel-Duby, Myocardin-related transcription factors are required for cardiac development and function. Dev. Biol. 406, 109–116 (2015).

8. Y. Guo, N. J. VanDusen, L. Zhang, W. Gu, I. Sethi, S. Guatimosim, Q. Ma, B. D. Jardin, Y. Ai, D. Zhang, B. Chen, A. Guo, G.-C. Yuan, L.-S. Song, W. T. Pu, Analysis of Cardiac Myocyte Maturation Using CASAAV, a Platform for Rapid Dissection of Cardiac Myocyte Gene Function In Vivo. Circ. Res. 120, 1874–1888 (2017).

9. Y. Guo, W. T. Pu, Genetic Mosaics for Greater Precision in Cardiovascular Research. Circ. Res. 123, 27–29 (2018).

10. A. C. H. Murphy, P. W. Young, The actinin family of actin cross-linking proteins - a genetic perspective. Cell Biosci. 5, 49 (2015).

11. N. J. VanDusen, Y. Guo, W. Gu, W. T. Pu, CASAAV: A CRISPR-Based Platform for Rapid Dissection of Gene Function *In Vivo*. Curr. Protoc. Mol. Biol. (available at http://onlinelibrary.wiley.com/doi/10.1002/cpmb.46/full).

12. S. Werfel, A. Jungmann, L. Lehmann, J. Ksienzyk, R. Bekeredjian, Z. Kaya, B. Leuchs, A. Nordheim, J. Backs, S. Engelhardt, H. A. Katus, O. J. Müller, Rapid and highly efficient inducible cardiac gene knockout in adult mice using AAV-mediated expression of Cre recombinase. Cardiovasc. Res. 104, 15–23 (2014).

13. T. W. Prendiville, H. Guo, Z. Lin, P. Zhou, S. M. Stevens, A. He, N. VanDusen, J. Chen, L. Zhong, D.-Z. Wang, G. Gao, W. T. Pu, Novel Roles of GATA4/6 in the Postnatal Heart Identified through Temporally Controlled, Cardiomyocyte-Specific Gene Inactivation by Adeno-Associated Virus Delivery of Cre Recombinase. PLoS One. 10, e0128105 (2015).

14. R. J. Platt, S. Chen, Y. Zhou, M. J. Yim, L. Swiech, H. R. Kempton, J. E. Dahlman, O. Parnas, T. M. Eisenhaure, M. Jovanovic, D. B. Graham, S. Jhunjhunwala, M. Heidenreich, R. J. Xavier, R. Langer, D. G. Anderson, N. Hacohen, A. Regev, G. Feng, P. A. Sharp, F. Zhang, CRISPR-Cas9 knockin mice for genome editing and cancer modeling. Cell. 159, 440–455 (2014).

15. B. Chen, C. Zhang, A. Guo, L.-S. Song, In situ single photon confocal imaging of cardiomyocyte T-tubule system from Langendorff-perfused hearts. Front. Physiol. 6, 134 (2015).

16. L. Madisen, T. A. Zwingman, S. M. Sunkin, S. W. Oh, H. A. Zariwala, H. Gu, L. L. Ng, R. D. Palmiter, M. J. Hawrylycz, A. R. Jones, E. S. Lein, H. Zeng, A robust and high-throughput Cre reporting and characterization system for the whole mouse brain. Nat. Neurosci. 13, 133–140 (2010).

17. D. O. Fürst, W. M. Obermann, P. F. van der Ven, Structure and assembly of the sarcomeric M band. Rev. Physiol. Biochem. Pharmacol. 138, 163–202 (1999).

18. A. Guo, L.-S. Song, AutoTT: automated detection and analysis of T-tubule architecture in cardiomyocytes. Biophys. J. 106, 2729–2736 (2014).

19. A. Subramanian, P. Tamayo, V. K. Mootha, S. Mukherjee, B. L. Ebert, M. A. Gillette, A. Paulovich, S. L. Pomeroy, T. R. Golub, E. S. Lander, J. P. Mesirov, Gene set enrichment analysis: A knowledge-based approach for interpreting genome-wide expression profiles. Proceedings of the National Academy of Sciences. 102(2005), pp. 15545–15550.

20. J. R. White, P. H. Naccache, R. I. Sha’afi,Stimulation by chemotactic factor of actin association with the cytoskeleton in rabbit neutrophils. Effects of calcium and cytochalasin B. J. Biol. Chem. 258, 14041–14047 (1983).

21. M. R. Bubb, I. Spector, A. D. Bershadsky, E. D. Korn, Swinholide A is a microfilament disrupting marine toxin that stabilizes actin dimers and severs actin filaments. J. Biol. Chem. 270, 3463–3466 (1995).

22. C. Y. Ho, D. E. Jaalouk, M. K. Vartiainen, J. Lammerding, Lamin A/C and emerin regulate MKL1-SRF activity by modulating actin dynamics. Nature. 497, 507–511 (2013).

23. M. A. Trembley, P. Quijada, E. Agullo-Pascual, K. M. Tylock, M. Colpan, R. A. Dirkx Jr, J. R. Myers, D. M. Mickelsen, K. de Mesy Bentley, E. Rothenberg, C. S. Moravec, J. D. Alexis, C. C. Gregorio, R. T. Dirksen, M. Delmar, E. M. Small, Mechanosensitive Gene Regulation by Myocardin-Related Transcription Factors Is Required for Cardiomyocyte Integrity in Load-Induced Ventricular Hypertrophy. Circulation. 138, 1864–1878 (2018).

24. A. Descot, R. Hoffmann, D. Shaposhnikov, M. Reschke, A. Ullrich, G. Posern, Negative regulation of the EGFR-MAPK cascade by actin-MAL-mediated Mig6/Errfi-1 induction. Mol. Cell. 35, 291–304 (2009).

25. B. Cen, A. Selvaraj, R. C. Burgess, J. K. Hitzler, Z. Ma, S. W. Morris, R. Prywes, Megakaryoblastic leukemia 1, a potent transcriptional coactivator for serum response factor (SRF), is required for serum induction of SRF target genes. Mol. Cell. Biol. 23, 6597–6608 (2003).

26. S. Y. Boateng, R. J. Belin, D. L. Geenen, K. B. Margulies, J. L. Martin, M. Hoshijima, P. P. de Tombe, B. Russell, Cardiac dysfunction and heart failure are associated with abnormalities in the subcellular distribution and amounts of oligomeric muscle LIM protein. Am. J. Physiol. Heart Circ. Physiol. 292, H259–69 (2007).

27. S. Arber, J. J. Hunter, J. Ross, M. Hongo, G. Sansig, J. Borg, J.-C. Perriard, K. R. Chien, P. Caroni, MLP-Deficient Mice Exhibit a Disruption of Cardiac Cytoarchitectural Organization, Dilated Cardiomyopathy, and Heart Failure. Cell. 88 (1997), pp. 393–403.

28. T. D. O’Connell, M. C. Rodrigo, P. C. Simpson, in Cardiovascular Proteomics, F. Vivanco, Ed. (Humana Press; http://link.springer.com/protocol/10.1385%2F1-59745-214-9%3A271), Methods in Molecular BiologyTM, pp. 271–296.

29. Y. Guo, Y. Kim, T. Shimi, R. D. Goldman, Y. Zheng, Concentration-dependent lamin assembly and its roles in the localization of other nuclear proteins. Mol. Biol. Cell. 25, 1287–1297 (2014).

30. Y. Guo, Y. Zheng, Lamins position the nuclear pores and centrosomes by modulating dynein. Mol. Biol. Cell. 26, 3379–3389 (2015).

31. S. Picelli, Å. K. Björklund, O. R. Faridani, S. Sagasser, G. Winberg, R. Sandberg, Smart-seq2 for sensitive full-length transcriptome profiling in single cells. Nat. Methods. 10, 1096–1098 (2013).

32. A. Dobin, C. A. Davis, F. Schlesinger, J. Drenkow, C. Zaleski, S. Jha, P. Batut, M. Chaisson, T. R. Gingeras, STAR: ultrafast universal RNA-seq aligner. Bioinformatics. 29, 15–21 (2013).

33. Y. Liao, G. K. Smyth, W. Shi, featureCounts: an efficient general purpose program for assigning sequence reads to genomic features. Bioinformatics. 30, 923–930 (2014).

34. M. I. Love, W. Huber, S. Anders, Moderated estimation of fold change and dispersion for RNA-seq data with DESeq2. Genome Biol. 15, 550 (2014).

35. A. Subramanian, P. Tamayo, V. K. Mootha, S. Mukherjee, B. L. Ebert, M. A. Gillette, A. Paulovich, S. L. Pomeroy, T. R. Golub, E. S. Lander, Others, Gene set enrichment http://nar.oxfordjournals.org/Downloaded from analysis: a knowledge-based approach for interpreting genome-wide expression profiles. Proc. Natl. Acad. Sci. U. S. A. 102, 15545–15550 (2005).

36. F. Ramírez, F. Dündar, S. Diehl, B. A. Grüning, T. Manke, deepTools: a flexible platform for exploring deep-sequencing data. Nucleic Acids Res. 42, W187–91 (2014).

37. J. T. Robinson, H. Thorvaldsdóttir, W. Winckler, M. Guttman, E. S. Lander, G. Getz, J. P. Mesirov, Integrative genomics viewer. Nat. Biotechnol. 29, 24–26 (2011).

